# Integrating electric field modelling and neuroimaging to explain inter-individual variability of tACS effects

**DOI:** 10.1101/581207

**Authors:** Florian H. Kasten, Katharina Duecker, Marike C. Maack, Arnd Meiser, Christoph S. Herrmann

## Abstract

Understanding variability of transcranial electrical stimulation (tES) effects is one of the major challenges in the brain stimulation community. Promising candidates to explain this variability are individual anatomy and the resulting differences of electric fields inside the brain. We integrated individual simulations of electric fields during tES with source-localization to predict variability of transcranial alternating current stimulation (tACS) aftereffects on α-oscillations. In two experiments, participants received 20 minutes of either α-tACS (1 mA) or sham stimulation. Magnetoencephalogram was recorded for 10 minutes before and after stimulation. tACS caused a larger power increase in the α-band as compared to sham. The variability of this effect was significantly predicted by measures derived from individual electric field modelling. Our results directly link electric field variability to variability of tACS outcomes, stressing the importance of individualizing stimulation protocols and providing a novel approach to analyze tACS effects in terms of dose-response relationships.

## 1 Introduction

Methods to non-invasively modulate brain activity via the transcranial application of magnetic or electrical stimulation are increasingly used in neuroscience to establish causal relationships between specific regions, or activation patterns (e.g. oscillations) in the brain and their behavioral correlates^1,2^. Among these techniques, transcranial electrical stimulation (tES) using weak direct (tDCS) or alternating (tACS) currents are of particular interest as they provide safe and tolerable stimulation at low costs and high portability^3,4^. These features render tES approaches promising for a wide range of clinical applications^5–7^. While tDCS is thought to exhibit its effect by changing neuronal excitability via tonic alterations of neuron’s resting membrane polarization^1,8–10^, the rhythmic shifts in the membrane potentials during tACS are believed to result in neuronal entrainment^2,11^. In addition, both methods have been reported to cause changes outlasting the duration of stimulation by several minutes to more than an hour^12–14^, likely via NMDA-receptor mediated plasticity^14–17^.

In recent years, tES methods received considerable criticism, arguing that stimulation effects are weak, highly variable or cannot be replicated^18–21^. Some authors even questioned whether current intensities in the range of 1 – 2 mA commonly used for tES cause sufficient electric field strengths inside the brain to elicit effects^22,23^. A variety of factors have been identified that can influence effects of non-invasive brain stimulation and may account for its variability^24–29^. A potential major source of tES variability are the influence of individual anatomy and the resulting differences of electric fields inside the brain^30,31^. The development of sophisticated computational models^32–34^ allows to study these differences using simulations. Recently, efforts have been carried out to validate the predictions of these models using in-vivo electrophysiological recordings in animals and humans^31,35,36^. Results from such simulated electric fields demonstrated that, when using a fixed stimulation montage and intensity, individual anatomical differences can cause substantial variability of electric fields inside the brain in terms of their spatial distribution and strength^30^. However, if and to which extent these differences explain variability of tES effects on behavioral or physiological outcome measures remains elusive.

In the current study, we investigated whether measures derived from individualized simulations of electric fields and source localization of the target brain activity can be used to explain variability of tACS effects. Specifically, we tested whether the spatial correlation of the target brain activity (spatial pattern of the source-projected α-oscillation) with the individually simulated electric field as well as the maximum field strength inside gray and white matter compartments can predict the variability of the power increase in the α-band after tACS. This power increase is relatively well established and has been repeatedly replicated^6,13,16,17,37^. The spatial correlation provides a measure of precision, namely how well does the electric field match the spatial pattern of the targeted brain activity, which is the source of the α-oscillation in the current study. The maximum field strength provides a measure of the intensity at which the target activity can be perturbed. In addition to the spatial precision of the stimulation, the precision of the stimulation frequency has to be considered when targeting brain oscillations using tACS. Recent work emphasized a possible role of the frequency relation between stimulation frequency and the frequency of the target oscillation in the generation of aftereffects^17,38^. While the frequency of the α-oscillation has long been assumed to be relatively stable, more recent evidence suggests that α-frequency can exhibit substantial intra-individual variability across different tasks and over time^39,40^. This mismatch was thus also included in our analysis. We hypothesized that a model incorporating these factors, which capture the quality of the targeting of stimulation, can explain variability of the power increase in the experimental group after receiving tACS, but not in a control group receiving sham stimulation. Our results indicate that a complex interplay between the spatial precision and strength of the electric field along with the mismatch of the stimulation frequency and participants’ individual α-frequency account for a large proportion of the variability of tACS aftereffects in humans.

## 2 Results

In the first experiment, a total of 40 volunteers received either 20-min of tACS or sham stimulation at their individual α-frequency (IAF), determined from a short, 3-min resting magnetoencephalogram (MEG) with eyes-open prior to the experiment. Their neuromagnetic activity with eyes-open in rest was recorded for 10-min immediately before and after stimulation (**Fig. 1a-c**). Based on a structural MRI of each subject, we performed an individual simulation of the expected electric field in the brain. Simulations were used to compute spatial correlations between electric fields and topographies of the α-source (IAF ± 2 Hz) during the pre-stimulation block obtained from a DICS (dynamic imaging of coherent sources) beamformer^41^. In addition, we extracted the average field strength among the 10,000 voxels with the highest field strength inside gray and white matter compartments (**Fig. 1d**).

**Figure 1:**
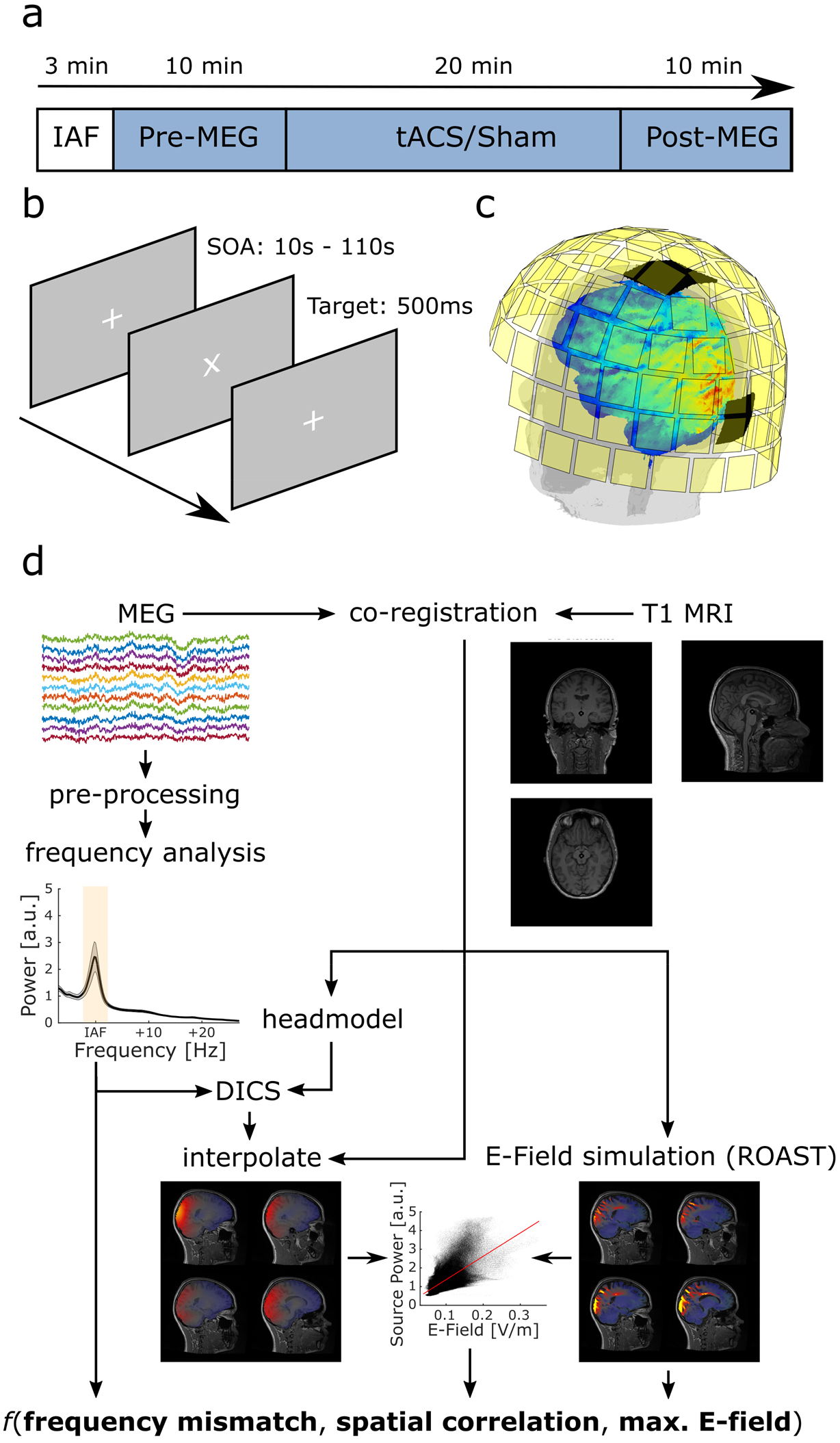
Experimental design and analysis pipeline. **(a)** Timecourse of the experiment. Prior to the main experiment, 3-min of eyesopen MEG were acquired to determine participants’ individual α-frequency (IAF). After 10-min of baseline measurement, participants received 20-min of tACS at IAF with 1 mA (peak-to-peak) or sham stimulation. Another 10-min of post-stimulation MEG were acquired thereafter. **(b)** Visual change detection task. Participants were instructed to detect rotations of a white fixation cross, presented on a screen at a distance of ~1m. **(c)** MEG sensor array and tACS montage. MEG was acquired from 102 magnetometer and 204 planar gradiometers. Stimulation electrodes were placed centered above positions Cz and Oz of the international 10-10 system. **(d)** Analysis pipeline to obtain spatial correlation between participants’ α-topography and electric field as well as the maximum electric field magnitude inside the gray and white matter compartments and mismatch between tACS frequency and the dominant frequency during the baseline block.

### 2.1 Inter-individual variability of electric fields

Although we administered and simulated tACS with a fixed intensity of 1 mA (peak-to-peak) and the same Cz-Oz electrode montage (**Fig. 1c**), simulations of electric fields revealed differences across subjects in terms of peak electric field magnitude arriving inside the cortex as well as the spatial distribution of electric fields (**Fig. 2**). To estimate the electric field strength arriving inside the cortex we averaged the electric field magnitude over the 10,000 voxels inside gray and white matter compartments exhibiting the largest electric field magnitude. On average, electric field strength was *M* = 0.13 V/m ± *SD* = 0.05 V/m (min = 0.08 V/m, max = 0.36 V/m). To characterize the similarity of electric fields across subjects, individual simulation results were warped into Montreal Neurological Institute (MNI)-space. Spatial correlations of the fields were computed between all subjects to attain insights into the overall variability of the factor. On average, electric fields correlated with *M* = .74 ± *SD* = .05 (*r*_*min*_ = .53, *r*_*max*_ = .85; **Fig. 2 bottom**). Spatial correlation between each subjects’ α-topography with the simulated electric field was on average *M* = .55 ± *SD* = .18 (*r*_*min*_ = −.12, *r_max_* = .76).

**Figure 2:**
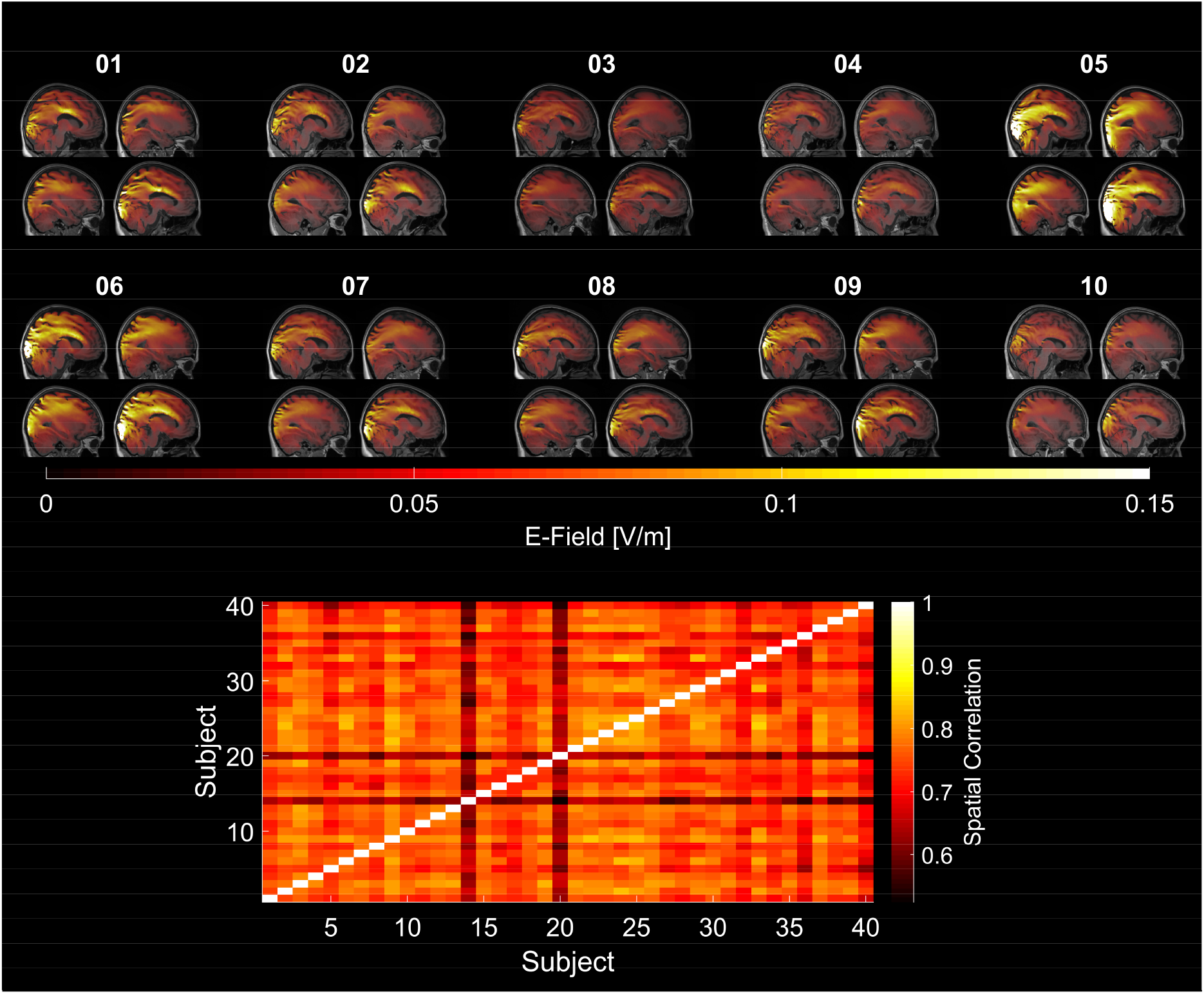
Variability of electric fields across subjects. **(Top)** Simulations of the electric fields inside the brain resulting from the Cz-Oz configuration applied at 1 mA (peak-to-peak) exemplified for the first ten subjects. Simulations were performed on the individual brain and warped into Montreal Neurological Institute (MNI) space for visualization purposes. Overall simulation results show a quite large variability between subjects. Please refer to Supplementary Fig. S1 for an overview of all simulations in the sample (**Bottom**) Spatial correlations of electric fields between all subjects in MNI-space.

### 2.2 Alpha power increase after tACS

In accordance with previous findings, a comparison of the source-projected power increase in the α-band from the pre- and post-stimulation blocks of the two experimental groups by means of an independent samples random permutation cluster t-test revealed a significantly larger power increase in the tACS group as compared to sham (*p*_*cluster*_ = .013; **Fig. 3**). No such effect was observed in the β- (all *p*_*cluster*_ > .18) or θ-range (all *p*_*cluster*_ > .19). The two groups did not differ with respect to their source-level α-power during the baseline block (*p*_*cluster*_ = 1). In both groups, dependent samples cluster permutation t-tests against baseline revealed a significant power increase in the α-range from the pre- to the post-stimulation block (tACS: *p*_*cluster*_ < .001; sham: *p*_*cluster*_ = .023; **Fig. 4a,f**). While the power increase in the sham group is limited to few occipital, posterior-parietal and temporal regions, the power increase in the tACS group spans a wide range of cortical areas covering occipital-parietal, temporal and frontal areas (**Fig. 4a**).

**Figure 3:**
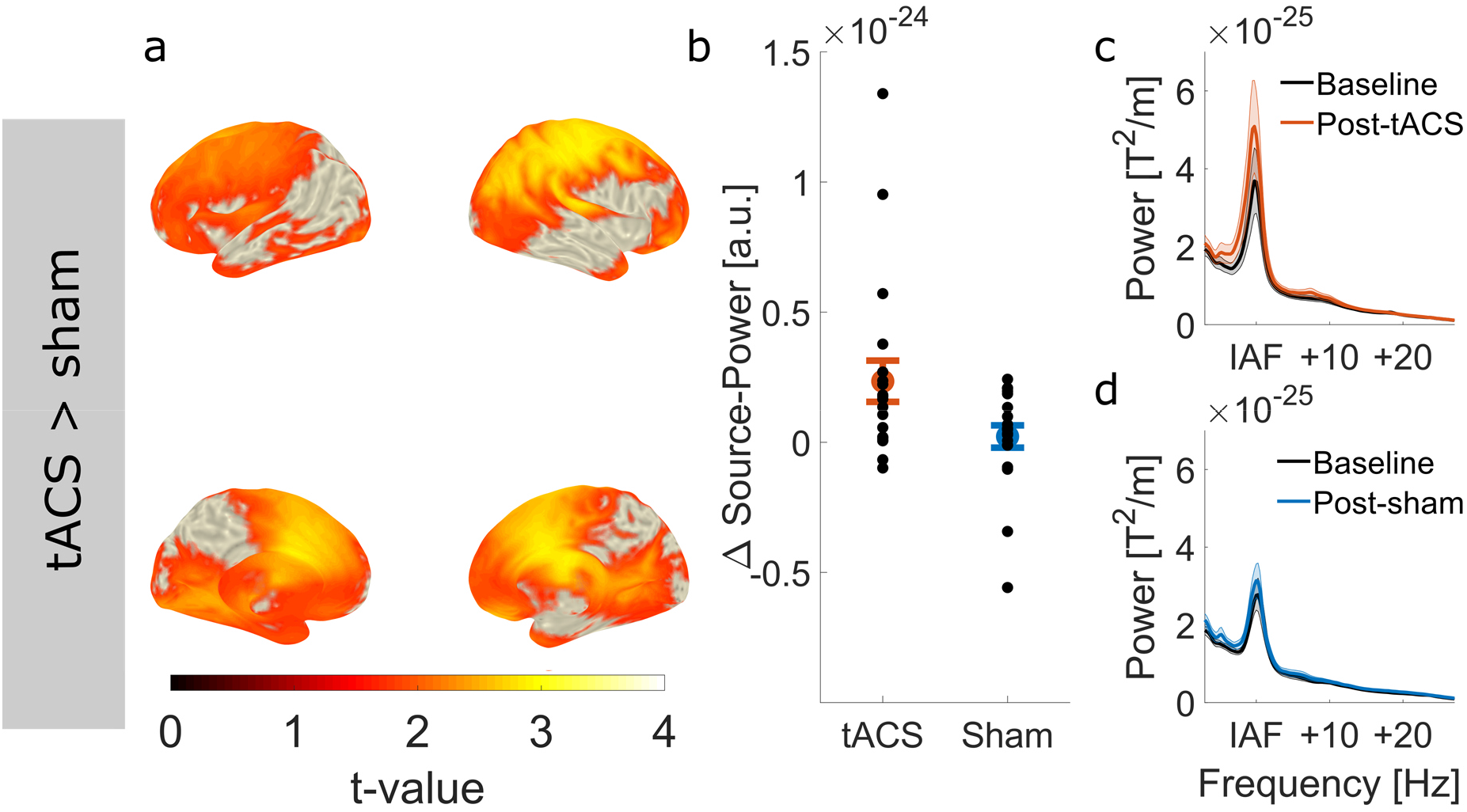
Effect of tACS on source-level α-power. **(a)** Statistical map contrasting the power increase from the pre- to the post-stimulation block between experimental conditions (tACS vs. sham). Statistical map shows t-values, thresholded at an α-level of .05. **(b)** Power increase within the cluster for each of the experimental groups. Error bars depict standard error of the mean (S.E.M.). Black dots represent the power increase of each individual subjects in the experimental groups. **(c)** Power spectra before and after tACS (average over all gradiometer sensors and participants). All spectra were aligned to participants’ individual α-frequency (IAF) before averaging. Shaded areas depict S.E.M. **(d)** Power spectra before and after sham stimulation.

**Figure 4:**
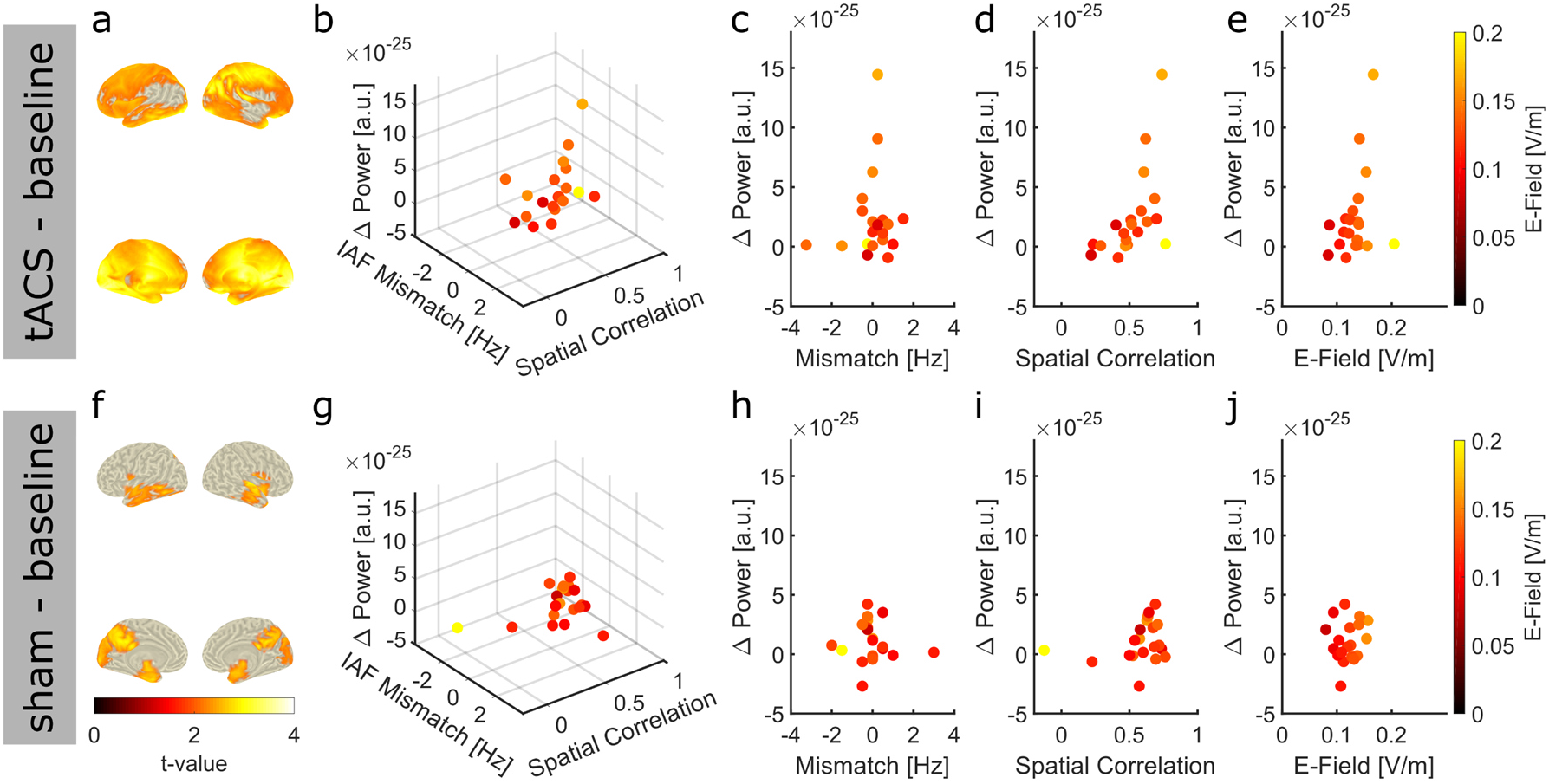
Power increase as a function of spatial correlation, field strength and frequency mismatch. **(a)** Statistical map contrasting post-tACS vs. pre-tACS power in the α-band. Statistical map depicts t-values thresholded at an α-level of .05. The cluster was used as ROI to extract the individual power increase of each subject in the tACS group for the subsequent regression analysis (**b-e**). (**b**) Power increase of the tACS group as a function of frequency mismatch between tACS frequency and the dominant frequency during baseline, the spatial correlation between the simulated electric field and the source-level α-topography during baseline, and the maximum field strength in gray and white matter compartments. Each dot represents data of a single subject. (**c-e**) Same data as in (**b**) shown for each predictor of the regression model. (**f**) Statistical map contrasting post-sham vs. pre-sham power in the α-band. Statistical map is thresholded at an α-level of .05. The cluster was used as ROI to extract the individual power increase of each subject in the sham group for the subsequent regression analysis. (**g**) Power increase of the sham group as a function of frequency mismatch between tACS frequency and the dominant frequency during baseline, the spatial correlation between the simulated electric field and the source-level α-topography during baseline, and the maximum field strength in gray and white matter compartments. Each dot represents data of a single subject. (**h-j**) Same data as in (**g**) shown for each predictor of the regression model.

### 2.3 Electric field variability predicts power increase after tACS

In order to evaluate whether the observed inter-individual differences of electric fields account for the variability of our outcome measure, for each subject, the average power increase between pre- and post-stimulation was extracted from the two group specific clusters, that is the cluster of each group exhibiting significant power increase from the pre- to the post-stimulation block **Fig. 4a,f.** The results were submitted to a multiple linear regression with factors *CONDITION* (tACS vs. sham), *PRECISION*_*spat*_ (spatial correlation of α-topography with electric field), *PRECISION*_*Freq*_ (mismatch between stimulation frequency and individual α-frequency during the baseline block) and *STRENGTH* (average over 10,000 highest electric field magnitudes inside gray and white matter). This full model was compared to all other possible models with subsets of the above factors using Akaike’s Information Criterion (AIC, **Supplementary Table S1**). The full model was retained for analysis as it exhibited the lowest AIC. Results of the regression analysis indicated that the four predictors explained 76 % of the variance (*R*^*2*^ = .76, *F*_*15,24*_ = 5.06, *p* < .001). More specifically, we found that the factor *CONDITION* (*β* = 2.51e-25, *t*_*24*_ = 2.38, *p* = .03), as well as interactions between *CONDITION*\**PRECISION*_*Freq*_*STRENGTH (*β* = 2.36e-23, *t*_*24*_ = 3.06, *p* = .005) and *CONDITION*\**PRECISION*_*Freq*_\**PRECISION*_*spat*_*STRENGTH (*β* = 1.56e-22, *t*_*24*_ = 3.47, *p* < .001) significantly predicted the power increase. In addition, there was a trend for an interaction of *CONDITION*\**PRECISION*_*spat*_\**PRECISION*_*Freq*_ (*β* = 2.36e-24, *t*_*24*_ = 2.05, *p* = .052).

All significant predictors explaining participants’ power increase involved the factor *CONDITION*. This pattern of results was expected given that our predictors are intended to relate to the efficacy of tACS and should thus not be suited to explain variance in the sham group. To specifically test that this is the case, we separately fitted a model with factors *PRECISION*_Spat_, *PRECISION*_Freq_ and *STRENGTH* to the data of the two experimental groups. In the tACS group, the model significantly predicts participants’ power increase (*R*^*2*^ = .87, *F*_*7,12*_ = 11.5, *p* < .001; **Fig. 4b-e, Fig. 5a**). The factors *PRECISION*_Spat_ (*β* = 1.68e-24, *t*_*12*_ = 4.26, *p* = .001) and *PRECISION*_*Freq*_ (*β* = 1.27e-25, *t*_*12*_ = 2.41, *p* = .003), as well as interactions of *PRECISION*_Spat_* *STRENGTH* (*β* = 2.64e-23, *t*_*12*_ = 2.99, *p* = .01), *PRECISION*_Spat_\**PRECISION*_Freq_*(*β* = 2.62e-24, *t*_*12*_ = 4.29, *p* = .001), *PRECISION*_Freq_* *STRENGTH* (*β* = 2.03e-23, *t*_*12*_ = 5.79, *p* < .001) and *PRECISION*_Spat_\**PRECISION*_Freq_* *STRENGTH* (*β* = 1.53e-22, *t*_*12*_ = 6.18, *p* < .001) significantly predicted participants’ power increase. Again, AIC suggests that this full model is superior to all other possible models with fewer predictors (**Supplementary Table S2**). As expected, the model fails to predict the power increase in the sham group (*R*^*2*^ = .13, *F*_*7,12*_ = 0.26, *p* = .96; **Fig. 4g-j, Fig. 5b**). In line with these results, the lowest AICs were obtained for an intercept model omitting all predictors related to stimulation, further confirming that the model is not suited to explain data of the sham group (**Supplementary Table S3**).

**Figure 5:**
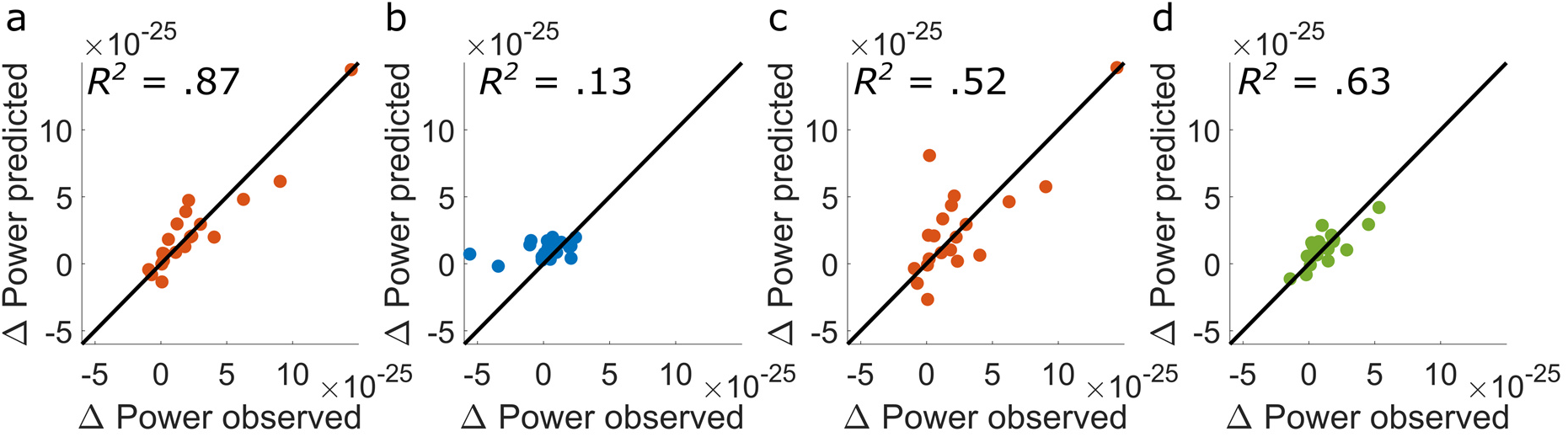
Model predictions. Scatterplots depict the power increase from the baseline to the poststimulation block predicted by the statistical models, plotted against empirically observed values. Each dot represents data of a single subject. The diagonals indicate the line of perfect prediction (predicted Δpower = observed Δpower). The distance of each datapoint from the diagonal indicates the residual prediction error. **(a)** Predicted and empirical power increase for the tACS group. **(b)** Predicted and empirical power increase in the sham group. **(c)** Power increase predicted for each subject in the tACS group by the cross-validated model. For each data point *n*_*i*_ the model was trained on *n*−*1* observations to predict the remaining *i*^*th*^ observation. **(d)** Individually sham controlled tACS effect (power increase relative to baseline after tACS – power increase relative to baseline after sham) on source-level α-power predicted by measures of tACS targeting.

Recently, concerns have been raised that tACS effects may not originate from electric stimulation of the brain, but exhibit its effects indirectly via stimulation of peripheral nerves (e.g. stimulation of the retina or transcutaneous nerves)^42,43^. Our results indicate that the extent of the tACS aftereffect can be predicted using the electric field inside the brain, which is difficult to explain with such peripheral mechanisms of action. We therefore conducted an additional analysis aiming to explain the data in our tACS group by a model incorporating the maximum current in the skin (*STRENGTHskin*; average over the maximum 10,000 voxels within the skin compartments) and the eyeballs (*STRENGTHeye*; average over the maximum 1000 voxels within the eyeballs). In addition, we included the factor *PRECISION*_*Freq*_ from our initial model as a similar effect of frequency mismatch has to be expected for peripheral stimulation effects. The resulting model was not able to significantly predict the power increase after tACS (*R*^*2*^ = .22, *F*_*7,12*_ = 0.49, *p* = .82). Based on AIC, no possible model incorporating a subset of these factors was superior to a simple intercept model in explaining the data (**Supplementary Table S4**). More importantly, none of the models was superior to the previous model incorporating the electric field in the brain (**Supplementary Table S3**, **S4**).

### 2.4 Model validation and replication

Because the model explains a striking amount of variance in the tACS group (~87%), we performed a leave one out cross-validation (LOOCV) to obtain a more conservative estimate of the explained variance. LOOCV can be used to perform cross-validation on small datasets. The model is trained (fitted) *n* times on *n*−1 datapoints and then used to predict the response variable for the remaining datapoint. This way, we can estimate how well the model generalizes to new observations. Based on the predictions of the LOOCV (**Fig. 5c**), we recomputed *R*^2^. Results suggest that the model still explains more than half (51.5%) of the variance in the tACS group (*R*^*2*^ = .52).

In order to investigate how much of participants’ individually sham controlled tACS effect the model can explain, and in order to replicate the previous results, we repeated the experiment using a within-subject design on a sample of 19 subjects. On two separate days, participants received tACS or sham stimulation for 20-min. The order of tACS and sham conditions were counterbalanced across participants.

Similar to the first experiment, a dependent samples random permutation cluster t-test revealed a stronger increase of participants’ source projected α-power after tACS as compared to sham (*p*_*cluster*_ < .001, **Fig. 6a-d**). Source projected α-power significantly increased after tACS (*p*_*cluster*_ < .001), and showed a trend towards increased α-power after sham stimulation (*p*_*cluster*_ = .08). Again, there was no significant difference in the neighboring β- (*p*_*cluster*_ > .16) and θ-frequency range (*p*_*cluster*_ > .08). Subsequently we tested whether participants’ individual stimulation effect in the α-band (power increase after tACS – power increase after sham) within the significant cluster (**Fig. 6a**) are explained by our measures of tACS targeting. To this end, the power increase relative to sham was submitted to a multiple linear regression with factors *PRECISION*_Spat_, *PRECISION*_Freq_ and *STRENGTH*. Although only reaching trend-level, the model again explains a striking amount of the variance of the tACS effect (*R*^*2*^ = .62, *F*_*7,11*_ = 2.66, *p* = .07; **Fig. 5d, Fig. 6e-h**). Specifically, *PRECISION*_Spat_ significantly predicted participants’ individual stimulation effect (*β* = 1.073e-24, *t*_*11*_ = 3.07, *p* = .01). In addition, there was a trend towards an interaction between *PRECISION*_Spat_, *PRECISION*_Freq_, and STRENGTH (*β* = −3.6e-23, *t*_*11*_ = −1.75, *p* = .1). Although the model performs weaker on the newly recorded dataset, results are generally in agreement with the findings of the previous experiment.

**Figure 6:**
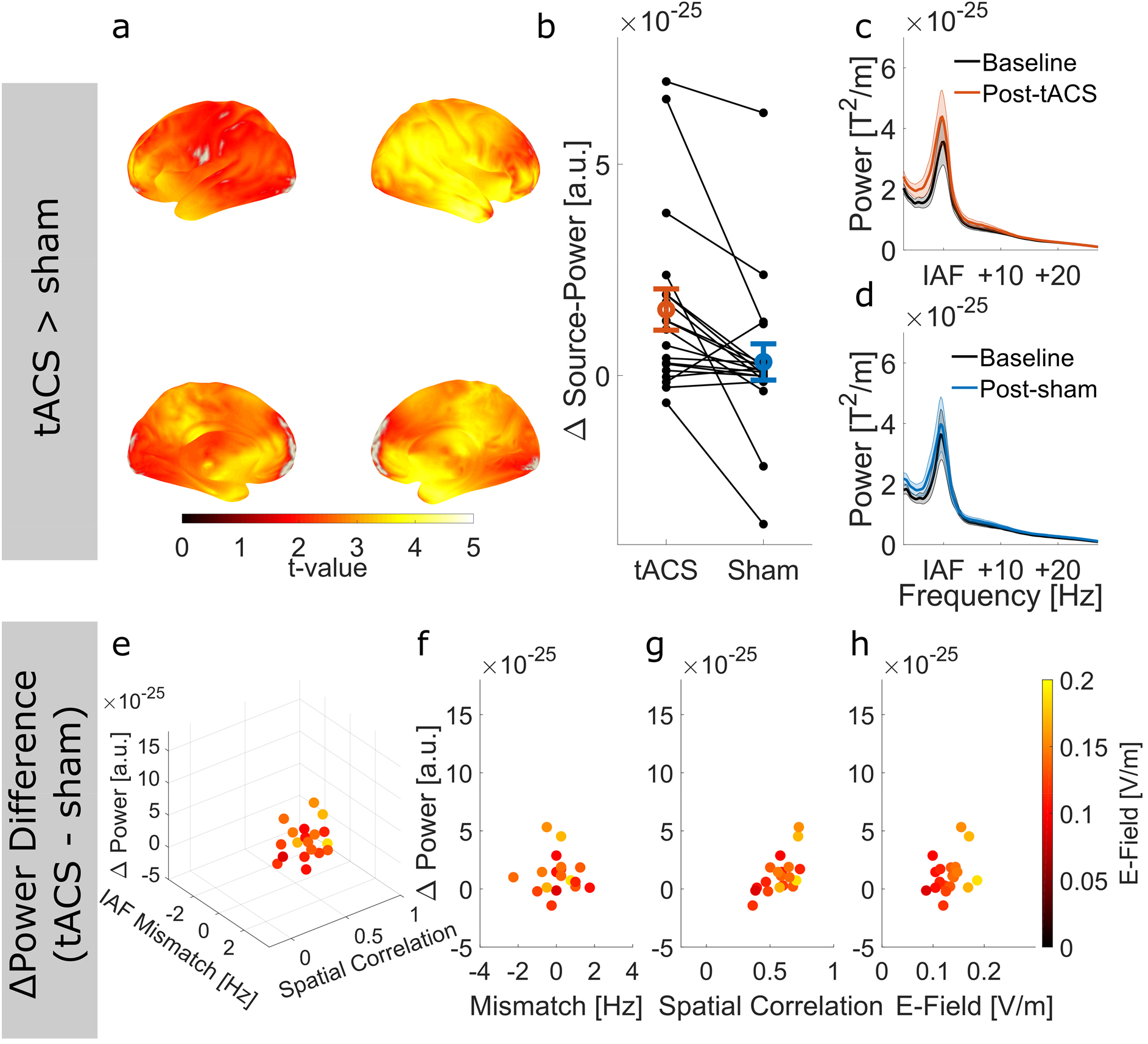
Results of within subject replication experiment. **(a)** Statistical map contrasting the power increase from the pre- to the post-stimulation block between experimental conditions (tACS vs. sham). Statistical map shows t-values, thresholded at an α-level of .05. **(b)** Power increase within the cluster for each of the experimental conditions. Error bars depict standard error of the mean (S.E.M.). Black dots/lines represent the power increase of each individual subjects in the experimental conditions. **(c)** Power spectra before and after tACS (average over all gradiometer sensors and participants). All spectra were aligned to participants’ individual α-frequency (IAF) before averaging. Shaded areas depict S.E.M. **(d)** Power spectra before and after sham stimulation. **(e)** Individually sham controlled tACS effect (power increase relative to baseline after tACS – power increase relative to baseline after sham) as a function of function of frequency mismatch between tACS frequency and the dominant frequency during baseline, the spatial correlation between the simulated electric field and the source-level α-topography during baseline, and the maximum field strength in gray and white matter compartments. Each dot represents data of a single subject. **(f-h)** Same data as in (**e**) shown for each predictor of the regression model.

## 3 Discussion

Increasing the reliability of low-intensity tES is one of the major challenges for the brain stimulation community. An understanding of the factors determining successful modulation of outcome measures by tES is crucial as the field is advancing these techniques towards clinical applications^5–7^. In the current study, we demonstrated that the variability of tACS aftereffects can be explained by an interplay of factors qualitatively capturing the targeting of the stimulation.

In line with previous findings^30^, our simulations indicate that electric fields induced at a fixed intensity with the same electrode montage vary quite substantially on an individual level. We were able to directly link this variability to the outcome of tACS. When integrated with neuroimaging, simulations of electric fields can be used to derive qualitative measures of the targeting (spatial correlation between electric field and target, maximum electric field strength inside the brain, mismatch between stimulation frequency and frequency of the target oscillation). Together these measures explained a substantial proportion of variance (~51% – 87%) of our outcome measure (power increase in the α-band after tACS). In contrast to this, the model did not explain any variance of the outcome measure after sham stimulation. In our second experiment, measures of tACS targeting explained about 63% of the variability of the individually sham-controlled stimulation effect in the sample. Taken together these results emphasize the importance of individualizing stimulation parameters for example by taking individual anatomy and the resulting electric field differences into account. Advancing algorithms for electric field modelling towards individualized electrode montages maximizing the field strength at the desired target^44^, and closed-loop stimulation systems adapting stimulation parameters to the current brain activity^45^ may greatly improve reliability of brain stimulation effects. This is especially important in clinical settings where the reliability of stimulation determines whether a patient’s symptoms improve.

In the context of research applications, study designs may benefit from incorporating individualized electric field modelling and neuroimaging for statistical analysis. We belief that this approach has some advantages over the pure comparison of group means, which is commonly used to investigate stimulation effects. Such comparisons implicitly assume that tES exerts consistent effects across participants. Especially when using “one-fits-all” stimulation protocols, a high prevalence of non-optimal targeting and the resulting numbers of potential low- or non-responders may compromise the sensitivity of such statistical approaches to detect stimulation effects.

In contrast, the statistical model proposed here tests for stimulation effects by assessing whether the variability of the outcome measure follows a dose-response relationship that would be expected based on the proposed underlying mechanisms. Consequently, the model is not only robust against low- and non-responders, but rather expects low- or non-responsiveness in cases where the standard stimulation protocol does not fit the individual subject well. As a further advantage, this mechanistic modelling largely rules out alternative explanations of the observed effects. In the field of tACS, concerns have been raised that stimulation effects could be explained by peripheral effects such as visual entrainment due to phosphenes^42^ (a perception of flickering lights resulting from a polarization of the retina^46^) or transcutaneous stimulation of peripheral nerves^43^. For the aftereffect observed in our study, it seems very unlikely that predictors derived from the electric field inside the brain would have been able to explain our data if such peripheral mechanisms had primarily caused the effect. This was further supported by our alternative model incorporating the electric fields in the skin and the eyeballs failing to predict participants’ power increase after tACS. To the contrary, as the model links the stimulation effect to variations of electric fields inside the brain results provide supporting evidence that tACS applied in the range of 1 mA can be sufficiently strong to elicit aftereffects arising from polarization of brain tissue. However, the impact of stimulation seems to depend on the strength and precision of the individual electric field and the precision of the stimulation frequency.

When the strength of tES-induced electric fields necessary to modulate neuronal activity is discussed, electric fields reaching the human brain are usually compared against thresholds derived from animal studies^11,47–49^. Those thresholds are in the range of 0.2 V/m to 0.5 V/m^47^. While evidence from animal models can strongly contribute to our understanding of the underlying mechanisms of tES methods, there are crucial discrepancies between experimental designs in animal and human studies that may limit the translation of voltage thresholds. In animal models, stimulation is usually applied to in-vitro brain slices or to localized neural assemblies via intracranial stimulation electrodes in-vivo. The modulation of neuronal activity is measured during short trains of stimulation, in the range of few seconds. In human studies, however, tES protocols commonly feature stimulation durations of several minutes (often >10-min), with stimulation applied to comparably large areas of the brain. Consequently, stimulation effects may build up over a longer periods of time or amplify via large-scale neuronal interactions^50^. Individual simulations suggest electric fields in our study were the range of 0.1 V/m – 0.2 V/m, providing evidence that the electric field strength necessary to elicit stimulation effects in humans may be in the lower range of those thresholds derived from animal models (or even below). More research will be necessary to determine the electric field strength required to modulate neuronal activity in humans (e.g. by testing stimulation protocols comparable to human experiments in animal models) and allow more informed discussions about tACS efficacy. Despite tuning tACS to each participants’ individual α-frequency as measured prior to the experiment, there was still a mismatch between the stimulation frequency and the individual α-frequency observed during the experiment that significantly contributed to the variability of the power increase after tACS. This mismatch has previously been reported to occur despite applying stimulation at participants’ individual frequency and to affect the extend tACS aftereffects^17,38^. Different processes may explain the occurrence of a frequency mismatch between tACS and brain oscillations. Firstly, the dominant frequency in a specific band may underly changes over time. For example, systematic drifts of the individual α-frequency have been observed over time and depending on the background task^39,40^. Secondly, for practical reasons in the current study the identified power peak in the α-band was rounded to the next integer frequency naturally giving rise to mismatches between stimulation frequency and the frequency of the targeted brain oscillation. Given the impact of this factor future studies might benefit from improved procedures to estimate tACS frequency.

Besides the investigation into the role of individual electric fields for tACS effects, the current study is among the first to perform source-localization of the tACS-induced power increase in the α-band. Although the effect has been repeatedly replicated, results usually rely on data from few electrode sites, providing little information about its spatial extend^13,16,17,37^. To our surprise, the effect of tACS in the α-band was very widespread covering a large proportion of the cortex, including frontal areas not covered by our electrode montage. We did not further investigate this observation up to this point as it was beyond the scope of our main research question. However, there is evidence that distributed brain networks communicate via correlated activity within specific frequency bands^51^. It might thus also be possible that the tACS-induced modulation of oscillatory activity within a circumscribed region could lead to co-stimulation of distant brain areas functionally coupled via the stimulated frequency band. It should however also be emphasized that differences in cluster extent are not independent of oscillatory power and might thus be solely explained by the power enhancement in the α-band. In addition to the power increase after tACS, we also observed an increase of power in the α-band after sham stimulation that was not explained by our statistical model. Such increase in α-power over time is commonly observed with time-on-task and has been associated with vigilance decrement and mental fatique^39,52–55^.

As with all scientific studies, some limitations of the current findings deserve consideration. The individual electric fields used for our analysis were obtained from computational modelling. This approach can only provide predictions of the individual electric field with an inherent degree of uncertainty and simplification. For example, errors in the automatic tissue segmentation can add random error to the estimated field strengths. Recently, first efforts have been carried out to validate and calibrate results of current flow predictions using in-vivo electrophysiology^31,35,36^. Results of these studies suggest that the models perform very well in predicting the spatial distribution of the induced electric fields, while tending to overestimate their strength. For our analysis approach, an accurate prediction of the exact field strength is not necessarily required, as long as the relative difference in the fields across subjects is accurately represented and uncertainties in the estimates, e.g. due to segmentation errors, introduce error variance but no systematic bias. Noteworthy, conductivities used for simulations of the ROAST toolbox have recently been calibrated to increase the accuracy of voltage and field strength predictions^35^. Our results indicate that both, the spatial distribution and the field strength predicted by individual electric field models, contain meaningful information allowing to predict the impact of tACS aftereffects, indicating that the computational models are sufficiently accurate to capture inter-individual differences. Nevertheless, further validation and optimization of electric field modelling using empirical data will be necessary to increase confidence in their predictions. Especially when models are integrated in the analysis of physiological or behavioral outcome measures as we propose in the current study, the accuracy of the utilized computational model will be crucial.

The predictions of the tACS aftereffect in the current study are based on predictors derived from the magnitude of the electric field inside the brain, ignoring the direction of the field relative to the cortical surface. While the current flow radial to the cortical surface (or normal component of the electric field) determines the strength of somatic polarization of cortical pyramidal cells, the current flow radial to the cortical surface polarizes horizontally arranged cortico-cortical axons^56^. In principle, these different components of the electric field could differentially contribute to stimulation effects. Models could incorporate this contribution by computing spatial correlations with the brain activity of interest and the strength parameter for each of the field directions. In the current experiment, we refrained from applying such models to the data as we aimed to keep statistical models sufficiently simple and interpretable.

Another important aspect to be discussed is the generalizability of the current results. Together with the mismatch of the stimulation frequency, individual differences of the electric fields explained a striking amount of the power increase in the α-band after tACS, pointing towards the significance of individual anatomy and the resulting differences in electric fields for tACS effects and potentially tES effects in general. Although the proposed underlying mechanisms of the different tES approaches differ^1,2^, they all ground on the principle that the electric fields induced to the brain alter the resting potential of neurons^8,11^, with stronger electric fields at the target area causing larger polarity changes. As this fundamental dose-response relationship is captured by our statistical model, it seems likely that individual differences in electric fields may have a similar impact for other tES methods or outcome measures. In the current study, we focused on the development of an analysis pipeline to investigate the impact of electric field differences on tES outcomes and tested it on a well replicated effect. Further work is needed to determine the exact impact of these differences for the various types of tES methods (tDCS, tACS, tRNS, etc.) and physiological and behavioral outcome measures, as well as for on- and offline effects. With the current work, we provide a powerful analysis framework, adaptable to EEG-source localization or fMRI that can strengthen our understanding of the contribution of individual anatomy on tES outcomes and the mechanisms of tES in general.

## 4 Online Methods

In experiment one, 40 healthy volunteers (age: 24 ± 3 years, 20 females, 20 males) without history of neurological or psychiatric disease were randomly assigned to one out of two experimental groups (tACS or sham) in a single-blind design. Groups were counterbalanced for participants’ sex. In experiment two, 22 healthy volunteers received tACS or sham stimulation on one out of two experimental sessions on separate days. The order of stimulation conditions was counterbalanced across participants. Both experimental sessions took place at the same time of day and were spaced at least four days apart. One participant aborted the experiment after the first session. Two participants indicated extreme levels of tiredness after the experiment as well as too short sleep durations the night prior to the experiment and were excluded from the study. Thus, 19 participants (11 females, age: 25 ± 3 years) remained for analysis. Two subjects participated in both experiments.

All subjects were right-handed according to the Edinburgh Handedness-Scale^57^ and had normal or corrected to normal vision. All were non-smokers and reported to be medication-free at the day of the measurement. Subjects gave written informed consent prior to the experiment and received monetary compensation for participation. Both experiments were approved by the *Commission for Research Impact assessment and Ethics* at the University of Oldenburg and performed in accordance with the declaration of Helsinki.

### 4.1 Magnetoencephalogram

Neuromagnetic signals were acquired at a rate of 1 kHz using a 306-channel whole-head MEG system with 102 magnetometer and 204 orthogonal planar gradiometer sensors (Elekta Neuromag Triux System, Elekta Oy, Helsinki, Finland), housed in an electrically and magnetically shielded room (MSR; Vacuumschmelze, Hanau, Germany). Five head-position indicator (HPI) coils were attached to participants’ head prior to the recording. Their positions were digitized along with the location of three anatomical landmarks (nasion, left and right tragus) and >200 head-shape samples using a Polhemus Fastrak (Polhemus, Colchester, VT, USA). Participants were seated underneath the sensor array in upright position (60° dewar orientation). To determine participants’ individual α-frequency (IAF), a three-minute recording of spontaneous MEG activity with eyes-open was acquired before the main experiment. Signals were filtered between 1 Hz and 40 Hz and segmented into 2-sec epochs. Fast Fourier Transforms (FFT; Hanning window) were computed for each of the segments and the resulting spectra were averaged. The power peak in the α-band between 8 Hz and 12 Hz within a fixed set of posterior gradiometer sensors (a detailed list is provided in the **Supplementary Materials**) was identified and the closest integer frequency to the identified peak was used as stimulation frequency during the following experiment. MEG was recorded with continuous head position tracking during two experimental blocks, one pre- and one post-stimulation (**Fig. 1a**). Although the recording was continued during stimulation, signals acquired during this period were discarded from the analysis due to the massive electromagnetic stimulation artifact, that can currently not be reliably removed from resting state recordings^58–61^.

### 4.2 Electrical Stimulation

Electrical stimulation was administered via two surface conductive rubber electrodes positioned centered over locations Cz (7 × 5 cm) and Oz (4 × 4 cm) of the international 10-10 system (**Fig. 1c**). Electrodes were attached to participants’ scalp using an electrically conductive, adhesive paste (ten20 paste, Weaver & Co, Aurora, CO, USA). The sinusoidal stimulation waveform was digitally sampled in Matlab 2016a at a rate of 10 kHz and streamed to a digital analog converter (Ni-USB 6251, National Instruments, Austin, TX, USA) connected to the remote input of a constant current stimulator (DC Stimulator Plus, Neuroconn, Illmenau, Germany). The stimulator was placed in an electrically shielded cabinet outside the MSR. From there, the signal was gated into the MSR via a tube in the wall using the MRI extension-kit of the stimulator (Neuroconn, Illmenau, Germany). Electrode impedance was kept below 20 kΩ (including two 5 kΩ resistors in the stimulator cables). Prior to the experiment, participants were introduced to potential sensations (visual and somatosensory) during stimulation and subsequently familiarized with the stimulation by brief application of tACS at the frequency and intensity used during the main experiment. Following a 10-min baseline period, participants received either 20-min of tACS at IAF or sham stimulation. Stimulation was applied with an intensity of 1 mA (peak-to-peak) and two 10-sec fade-in/fade-out intervals at the beginning and end of the stimulation period, respectively. During sham stimulation, tACS was applied during the first 30-sec of the stimulation period (fade-in and fade-out). All other stimulation parameters were kept similar. Stimulation frequency was on average *M* = 10.1 Hz ± *SD* = 1 Hz (*M*_*tACS*_ = 9.9 Hz ± 1 Hz; *M_Sham_* = 10.4 Hz ± 0.6 Hz) for the first experiment and 10.5 Hz ± 1.1 Hz (*M*_*tACS*_ = 10.5 Hz ± 1.1 Hz; *M*_*Sham*_ = 10.5 Hz ± 1.1 Hz) for the second experiment.

After the recordings, participants filled out a questionnaire assessing common adverse effects of transcranial electrical stimulation^62^ and indicated whether they believe they received tACS or sham stimulation. Subsequently, participants were informed about their true experimental condition and the goals of the study. Results of the debriefing are presented in the **Supplementary Materials**

### 4.3 Vigilance task

To ensure that participants remained awake and attentive during the 40-min measurement, they performed a visual change detection task similar to previous studies^13,16,38,63^ (**Fig. 1b**). Visual stimuli were presented with Matlab 2016a, using Psychtoolbox 3^64^. Stimuli were rear-projected onto a screen inside the MSR at a distance of ~100 cm. At the center of the screen a white fixation cross on a gray background was presented. The fixation cross was rotated by 45° for a duration of 500 ms at random intervals with an SOA of 10-sec – 110-sec (**Fig. 1b**). Participants were asked to react to the rotation by pressing a button with their right index finger.

### 4.4 MRI acquisition

To perform source analysis and individual simulations of electric fields during stimulation, a structural MRI was obtained from each subject. Images were acquired using a Siemens Magnetom Prisma 3T whole-body MRI machine (Siemens, Erlangen, Germany). A T1-weighted 3-D sequence (MPRAGE, TR = 2000 ms, TE = 2.07 ms) with a slice thickness of 0.75 mm was used.

### 4.5 Data Analysis

Data analysis was performed in Matlab 2016a (The MathWorks, Inc. Natick, MA, USA) using the Fieldtrip toolbox^65^ for MEG data processing and ROAST v. 2.7^34^ for individualized electric field modelling. Statistical analysis of source level data was performed using statistical functions provided by the Fieldtrip toolbox. All other statistical analyses were performed using R 3.5.1 (The R Core Team, R Foundation for Statistical Computing, Vienna, Austria).

#### 4.5.1 MEG preprocessing

External interference in the MEG was suppressed using the spatiotemporal signal space separation method (tSSS), with standard settings (L_in_ = 8, L_out_ = 3, correlation limit = .98) ^66,67^ using MaxFilter^TM^ (Elekta Neuromag, Elekta Oy, Finland). Movement compensation was performed using the continuous HPI signals^68^. The tSSS method decomposes the MEG signal into spatiotemporal components originating from inside and outside the sensor helmet. The method is commonly used to suppress external artifacts and interference signals, especially those originating in the proximity of the head (e.g. implants, deep-brain stimulator, etc.)^66,67,69^. The method is thus well suited to remove interference brought into the MEG helmet via the cables connected to the stimulation electrodes. Further, it allows to compensate for head-movements by transforming the signals to the initial head position^68^. Signals were subsequently imported to Matlab and resampled to 250 Hz. A 4^th^-order forward-backward Butterworth filter introducing zero phase-shift between 1 Hz and 40 Hz was applied. Artifacts reflecting heart-beat, eye-movements or muscle activity were manually removed using independent component analysis (ICA). After visual inspection of component topographies and time-courses an average of 3.7 (± SD: 1; min: 3, max: 8) components were removed before back-projecting the signals into sensor-space. In experiment 2, 3.6 (± SD: 0.9; min: 2, max: 6) components were rejected on average. Rejection criteria were based on recommendations in the literature^70^. Signals were cut into 2-sec epochs. Segments still containing artifacts were rejected. FFTs (4-sec zero-padding, Hanning window) were computed on each of the segments. The resulting power spectra were averaged across the first 260 artifact-free segments in each experimental block.

#### 4.5.2 DICS beamforming

Power in the individual α-band (IAF ± 2Hz) was projected into source-space using a DICS (dynamic imaging of coherent sources) beamformer^41^ utilizing all 306 (magnetometer and gradiometer) channels. A common spatial filter was computed from the averaged cross-spectrum in the IAF band across all segments of the two experimental blocks. Data were projected onto an equally spaced 6 mm grid, warped into MNI (Montreal Neurologic Institute) space. Single-shell head-models^71^, derived from individual T1-weighted MRIs, co-registered to participants’ head position inside the MEG were used. Regularization was set to λ = 1e-12. The common filters were then applied to project data of the pre- and post-stimulation block. For each source location, the power difference between the pre- and post-stimulation block was computed. To test whether the power increase in the α-band was larger after tACS as compared to sham, power differences were submitted to a one-sided non-parametric random permutation cluster t-test with 10,000 randomizations and Monte Carlo estimates to calculate p-values. In addition, power in the α-band before and after stimulation was compared separately for both groups using random permutation cluster t-tests for dependent samples. The identified clusters were used as region of interests (ROI) to extract the average power increase from the pre- to the post-stimulation block for subsequent analysis (see next section). To evaluate frequency specificity of the effects, the analysis was repeated for the individual theta (IAF − 5 Hz to IAF − 3 Hz) and beta (IAF + 4 to IAF + 20 Hz) band.

#### 4.5.3 Individualized electric field calculations

Individual simulations of the electric field induced by the Cz – Oz montage were performed on the co-registered, T1-weighted MRI of each subject using the ROAST toolbox v2.7^34^. The toolbox offers some advantages over other modelling tools currently available, as it requires comparably short computation times (~25-min per subject), automatically determines standard EEG electrode positions in individual head space and provides results in Matlab format, allowing easy integration with source-level MEG results. As part of the ROAST pipeline, a 6-compartment (white matter, gray matter, csf, bone, skin, air), finite-element model is created from individual MRIs using the SPM12 segmentation algorithm. A post-processing routine is subsequently used to optimize the segmentation output for electric field modelling (for details see^34^). Simulations were run with an injected current of 0.5 mA (corresponding to 1 mA peak-to-peak), a 7 × 5 cm rectangular electrode patch over electrode site Cz, and a 4 × 4 cm rectangular electrode patch over electrode site Oz. Electrodes were modelled with a thickness of 2 mm and a 2 mm layer of gel. Default conductivities of the toolbox were used for the different compartments. Recently, these have been validated/calibrated based on intracranial recordings of 10 human epilepsy patients^35^.

To capture the inter-individual variability of the electric fields across subjects, we computed two measures, one indicating the spatial precision of stimulation (*PRECISION*_*spat*_: how well does the electric field overlap with the targeted brain activity), the other indicating the strength of stimulation inside the brain (*STRENGTH*). As a measure of precision, we calculated the spatial correlation between the electric field with the individual topography of each participant’s α-power. To this end, participants’ IAF band power (IAF ± 2 Hz) during the pre-stimulation block was localized using a DICS beamformer. Data were projected onto an equally-spaced 3 mm grid defined in individual head-space (no warping onto a standard brain). Filters were computed using the cross-spectra in the IAF band obtained from the artifact free segments of the baseline block. To account for the center-of-head bias of the beamformer, the neural activity index (NAI) was computed. The NAI is the source activation at each dipole location normalized by an estimate of the noise at that location. The NAIs were subsequently interpolated onto the individual, T1-weighted MRI, which has the same resolution as the electric field calculation and thus allowed us to compute the spatial correlation between the source-projected α-topography and the individual electric field profile for each subject. To index the *STRENGTH* of stimulation inside the brain, we identified the 10,000 voxels inside the grey and white matter compartments of each subjects’ simulation result showing highest electric field magnitude and computed the average electric field magnitude across these voxels.

To evaluate whether these measures of electric field differences account for the variability of our outcome measure, we modelled each subject’s power increase within the ROIs as a function of *CONDITION* (tACS vs. sham), *PRECISION*_*spat*_, *STRENGTH* and *PRECISION*_*Freq*_ with a multiple linear regression model. *PRECISION*_*Freq*_ captures the mismatch between the pre-determined stimulation frequency (sf) and the dominant frequency (df) observed during the baseline block (mismatch = sf − df), extracted from the average spectrum over all sensors.

## 5 Data Availability

The data that support the findings of this study are available upon reasonable request from the corresponding author CSH. The data are not publicly available due to potentially identifying information that could compromise participant privacy. The source data underlying Fig. 2 (bottom), Fig. 3b-d, Fig. 4,5 and Fig. 6b-h are provided as a Source Data file.

## 6 Acknowledgements

The authors thank Helge Ahrens, Dr. Tina Schmitt, Gülsen Yanc and Katharina Grote for the acquisition of structural MRIs and for supporting the MEG data collection.

This research was supported by the Neuroimaging Unit of the Carl von Ossietzky University Oldenburg funded by grants of from the German Research Foundation (3T MRI INST 184/152-1 FUGG and MEG INST 184/148-1 FUGG). Christoph S. Herrmann was supported by a grant of the German Research Foundation (SPP, 1665 HE 3353/8-2).

## 7 Author Contributions

FHK and CSH conceived the study; FHK and KD programmed the experiment; FHK, KD and AM acquired data for experiment 1, FHK and MCM acquired data for experiment 2, FHK analyzed the data, all authors wrote the manuscript.

## 8 Conflict of interest

CSH holds a patent on brain stimulation and received honoraria as editor from Elsevier Publishers, Amsterdam. FHK, KD, MCM, and AM, declare no competing interests.

## 10 Supplementary Results

**Supplementary Fig. S1:**
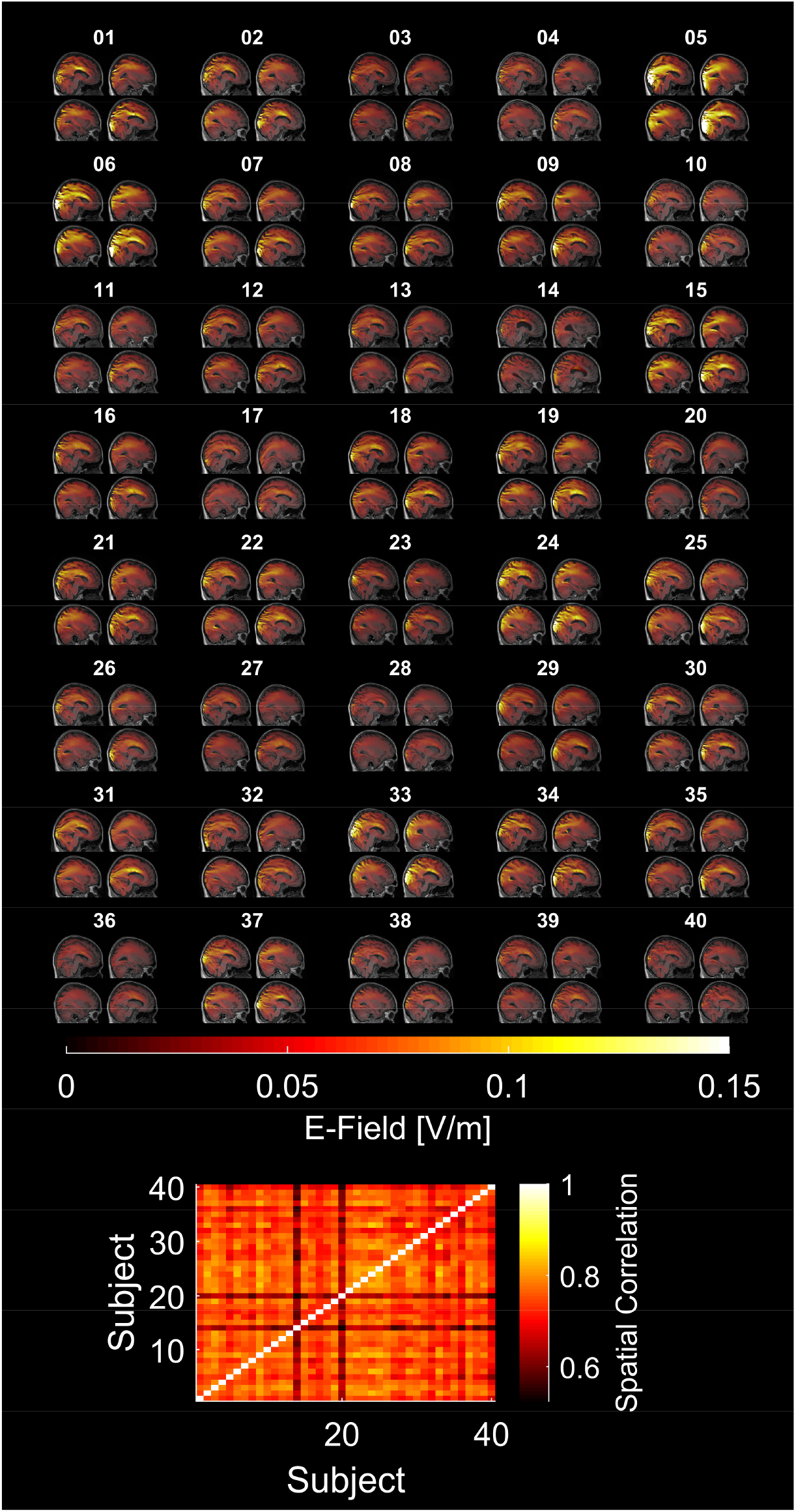
Variability of electric fields across all subjects. **(Top)** Simulations of the electric fields inside the brain resulting from the Cz-Oz configuration applied at 1 mA (peak-to-peak) shown for all subjects. Simulations were performed on the individual brain and warped into MNI space for visualization purposes. Overall simulation results show a quite large variability between subjects. Please refer to Supplementary Fig. S1 for an overview of all simulations in the sample (**Bottom**) Spatial correlations of electric fields between all subjects in MNI-space.

### 10.1 Model comparisons using Akaike’s Information Criterion

**Supplementary Table S1:** Comparison of Akaike’s Information Criterion (AIC) across all possible models for the first regression analysis. AIC clearly favors the full model (top).

| Model | AIC |
| --- | --- |
| *CONDITION\*PRECISION<sub>Spat</sub>\*PRECISION<sub>Freq</sub>\*STRENGTH* (final model) | -4429.8 |
| *CONDITION\* PRECISION<sub>Spat</sub>\*STRENGTH* | -4404.8 |
| *CONDITION\* PRECISION<sub>Spat</sub>\* PRECISION<sub>Freq</sub>* | -4405.2 |
| *CONDITION\* PRECISION<sub>Freq</sub>\*STRENGTH* | -4403.5 |
| *CONDITION\* STRENGTH* | -4403.1 |
| *CONDITION\* PRECISION<sub>Freq</sub>* | -4399.7 |
| *CONDITION\* PRECISION<sub>Spat</sub>* | -4412.1 |
| *CONDITION* | -4403.2 |
| *PRECISION<sub>Spat</sub>\*PRECISION<sub>Freq</sub>\*STRENGTH* | -4405.1 |
| *PRECISION<sub>Spat</sub>\* STRENGTH* | -4405.1 |
| *PRECISION<sub>Spat</sub>\*PRECISION<sub>Freq</sub>* | -4402.7 |
| *PRECISION<sub>Freq</sub>\* STRENGTH* | -4405.2 |
| *STRENGTH* | -4401.2 |
| *PRECISION<sub>Freq</sub>* | -4400.9 |
| *PRECISION<sub>Spat</sub>* | -4405.7 |
| *Intercept* | -4402.8 |

**Supplementary Table S2:** Comparison of Akaike’s Information Criterion (AIC) across all possible models for the second regression analysis fitted to data of the tACS group. Again, AIC clearly favors the full model (top).

| Model | AIC |
| --- | --- |
| $PRECISION_{Spat} * PRECISION_{Freq} * STRENGTH$ (final model) | -2217.3 |
| $PRECISION_{Spat} * STRENGTH$ | -2192.3 |
| $PRECISION_{Spat} * PRECISION_{Freq}$ | -2192.6 |
| $PRECISION_{Freq} * STRENGTH$ | -2192.1 |
| $STRENGTH$ | -2190.9 |
| $PRECISION_{Freq}$ | -2188.8 |
| $PRECISION_{Spat}$ | -2196.1 |
| Intercept | -2190.5 |

**Supplementary Table S3:** Comparison of Akaike’s Information Criterion (AIC) across all possible models for the second regression analysis fitted to data of the sham group. Here, AIC favors the intercept model omitting all other predictors. To allow direct comparisons of the model fits between the groups, the full model (top) was fitted to the data of the sham group.

| Model | AIC |
| --- | --- |
| $PRECISION_{Spat} * PRECISION_{Freq} * STRENGTH$ (final model) | -2211.0 |
| $PRECISION_{Spat} * STRENGTH$ | -2218.5 |
| $PRECISION_{Spat} * PRECISION_{Freq}$ | -2218.4 |
| $PRECISION_{Freq} * STRENGTH$ | -2216.5 |
| $STRENGTH$ | -2220.2 |
| $PRECISION_{Freq}$ | -2220.2 |
| $PRECISION_{Spat}$ | -2221.5 |
| Intercept | -2222.1 |

**Supplementary Table S4:** Comparison of Akaike’s Information Criterion (AIC) across all possible models for the alternative model incorporating the strength of peripheral stimulation of the skin and the eye-balls. The model was fitted to data of the tACS group. AIC favors the intercept model omitting all other predictors.

| Model | AIC |
| --- | --- |
| $PRECISION_{Freq} * STRENGTH_{Eye} * STRENGTH_{Skin}$ (full model) | -2181.5 |
| $PRECISION_{Freq} * STRENGTH_{Eye}$ | -2186.7 |
| $PRECISION_{Freq} * STRENGTH_{Skin}$ | -2185.5 |
| $STRENGTH_{Eye} * STRENGTH_{Skin}$ | -2187.8 |
| $STRENGTH_{Eye}$ | -2190.4 |
| $STRENGTH_{Skin}$ | -2189.0 |
| $PRECISION_{Freq}$ | -2188.8 |
| Intercept (best model) | -2190.5 |

### 10.2 List of MEG channels used to determine participants’ individual α-frequency

‘MEG1632’, ‘MEG1633’, ‘MEG1732’, ‘MEG1733’, ‘MEG1742’, ‘MEG1743’, ‘MEG1842’, ‘MEG1843’, ‘MEG1912’, ‘MEG1913’, ‘MEG1922’, ‘MEG1923’, ‘MEG1932’, ‘MEG1933’, ‘MEG1942’, ‘MEG1943’, ‘MEG2012’, ‘MEG2013’, ‘MEG2022’, ‘MEG2023’, ‘MEG2032’, ‘MEG2033’, ‘MEG2042’, ‘MEG2043’, ‘MEG2112’, ‘MEG2113’, ‘MEG2122’, ‘MEG2123’, ‘MEG2132’, ‘MEG2133’, ‘MEG2232’, ‘MEG2233’, ‘MEG2312’, ‘MEG2313’, ‘MEG2322’, ‘MEG2323’, ‘MEG2332’, ‘MEG2333’, ‘MEG2342’, ‘MEG2343’, ‘MEG2432’, ‘MEG2433’, ‘MEG2442’, ‘MEG2443’, ‘MEG2512’, ‘MEG2513’, ‘MEG2542’, ‘MEG2543’

### 10.3 Debriefing

#### Experiment 1

Overall 16 out of 40 participants indicated that they believed to have received electrical stimulation during the experiment (11 out of 20 in the sham group and 5 out of 20 in the tACS group). A Pearson’s Chi-Squared test for count data revealed no significant difference for the number of ‘yes’ and ‘no’ answers between groups (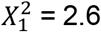, *p* = .11). In addition, we asked participants to rate their confidence in the answer on a scale from 1 to 10. Participants’ ratings were submitted to a 2×2 factorial Analysis of Variance (ANOVA) with between subject factors CONDITION (tACS vs. sham) and ANSWER (yes vs. no). The ANOVA revealed neither an effect of CONDITION (*F*_*1,36*_ = 2.00, *p* = .16, *η*^*2*^ = 0.05) or ANSWER (*F*_*1,36*_ = 0.007, *p* = .92, *η^2^* < 0.01), nor an interaction (*F*_*1,36*_ = 0.04, *p* = .54, *η*^*2*^ = 0.01). On average participant’s confidence was in the upper-medium range of the rating scale (tACS+yes: 7 ± 0.7, tACS+no: 6.33 ± 2.58, sham+yes: 5.18 ± 2.89, sham+no: 5.56 ± 2.31).

#### Experiment 2

One participant did not completely fill out the debriefing questionnaire and had to be excluded from the debriefing analysis. Of the remaining 18 participants 11 correctly indicated that they thought they had received tACS after the tACS session, while 7 indicated to believe not to have received active stimulation. After sham stimulation 12 participants indicated they thought to have received tACS, while 6 indicated that they did not thought to have received stimulation. A Pearson’s Chi-Squared test for count data revealed no significant difference for the number of ‘yes’ and ‘no’ answers between experimental sessions (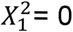, *p* = 1). There was no effect of CONDITION (*F*_*1,32*_ = 2.54, *p* = .12, *η*^*2*^ = 0.07), or ANSWER (*F*_*1,32*_ = 0.39, *p* = .53, *η*^*2*^ = 0.01) and no interaction effect (*F*_*1,32*_ = 0.04, *p* = .85, *η*^*2*^ < 0.01) on participants’ confidence ratings. On average participants’ confidence was in the upper medium range (tACS+yes: 5.36 ± 2.2, tACS+no: 6 ± 2.2, sham+yes: 6.7 ± 2.1, sham+no: 7 ± 2.6).

Overall, results of the debriefing indicate that participants in both experiments were successfully blinded towards their experimental condition.

### 10.4 Voxel wise correlation between electric field and tACS effect

**Supplementary Fig. S2:**
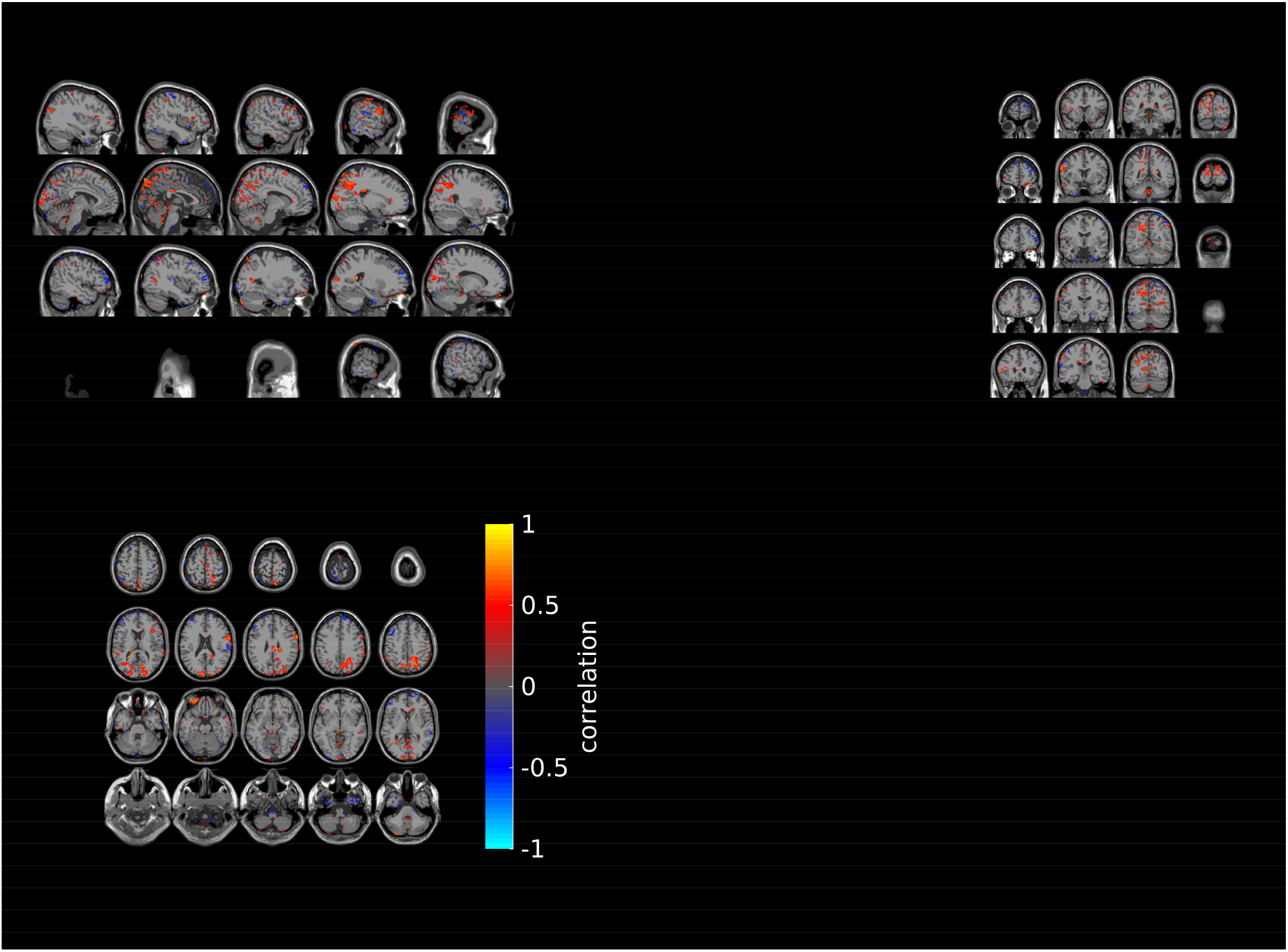
Voxel-wise correlation between electric field magnitude and tACS effect. Correlation between the simulated electric field and the individually sham controlled tACS effect in the α-band (α-power increase relative to baseline after tACS – α-power increase relative to baseline after sham). Pearson’s correlation coefficient between the simulated electric field warped into MNI space and the tACS effect was computed across subjects for each voxel. The resulting correlation maps are thresholded at a significance level of p < .05 (uncorrected).

### 10.5 Predicting the maximum power within the group specific cluster

Using a group ROI can introduce a bias such that larger power values are observed for participants whose α-power distribution on the source level is more similar to the cluster. We thus repeated our analysis identifying and averaging over the 1000 source locations within the clusters that show the strongest α-power increase to baseline for each participant. We submitted these power values to our linear regression analysis with factors CONDITION, PRECISION_Freq_, PRECISION_Spat_, and STRENGHT. We obtained similar results as for our ROI analysis in section 2.3. The model significantly predicted participants peak power increase in the ROI (*R*^*2*^ = .78, *F*_*15,24*_ = 5.89, *p* < .001). Specifically, the factors CONDITION (*β* = 7.203e-25, *t*_*24*_ = 3.33, *p* = .003), the interaction between CONDITION, PRECISION_Freq_ and STRENGTH (*β* = 5.114e-23, *t*_*24*_ = 3.24, *p* = .003) and the interaction between CONDITION, PRECISION_Freq_, PRECISION_Spat_ and STRENGTH (*β* = 2.896-22, *t*_*24*_ = 4.07, *p* < .001) significantly predicted the power increase. When separately fitted to the data of the two groups, the model again failed to explain the power increase in the sham group (*R*^*2*^ = .15, *F*_*7,12*_ = 0.31, *p* = .93), but significantly predicts the power increase after tACS (*R*^*2*^ = .82, *F*_*7,12*_ = 7.58, *p* = .001). Specifically, the factors PRECISION_Spat_ (*β* = 3.90-24, *t*_*12*_ = 3.74, *p* = .003), the interactions between PRECISION_Spat_ and PRECISION_Freq_ (*β* = 4.41-24, *t*_*12*_ = 2.73, *p* = .018), STRENGTH and PRECISION_Freq_ (*β* = 4.53-23, *t*_*12*_ = 4.89, *p* < .001) and PRECISION_Freq_, PRECISION_Spat_ and STRENGTH (*β* = 2.90-22, *t*_*12*_ = 6.56, *p* < .001) significantly predicted participants peak power increase after tACS.

### 10.6 Frequency mismatch for each subject

**Supplementary Table S5:** Overview of individual α-frequency measured before the experiment, stimulation frequency during the experiment, IAF measured during the baseline block and the mismatch between the tACS frequency and the IAF during the baseline block of the first experiment.

| ID | IAF before experiment | tACS frequency | IAF during baseline | Mismatch<br>tACS/baseline |
| --- | --- | --- | --- | --- |
| 01 | 11.5 | 11 | 11.25 | -0.25 |
| 02 | 10 | 19 | 10 | 0 |
| 03 | 11 | 11 | 11 | 0 |
| 04 | 11 | 11 | 11.5 | -0.5 |
| 05 | 10 | 10 | 11.5 | -1.5 |
| 06 | 11 | 11 | 11.25 | -0.25 |
| 07 | 10 | 10 | 10.5 | -0.5 |
| 08 | 8 | 8 | 11.25 | -3.25 |
| 09 | 9.5 | 10 | 11.5 | -1.5 |
| 10 | 9.5 | 10 | 10.25 | -0.25 |
| 11 | 10.5 | 10 | 8.5 | 1.5 |
| 12 | 10.5 | 10 | 10.5 | -0.5 |
| 13 | 10 | 10 | 9.5 | 0.5 |
| 14 | 12 | 12 | 11.5 | 0.5 |
| 15 | 10 | 10 | 10 | 0 |
| 16 | 11 | 11 | 10.25 | 0.75 |
| 17 | 10 | 10 | 10.5 | -0.5 |
| 18 | 8.5 | 8 | 8 | 0 |
| 19 | 10.5 | 11 | 10.5 | 0.5 |
| 20 | 11.5 | 12 | 11 | 1 |
| 21 | 11 | 9 | 11 | -2 |
| 22 | 9.5 | 10 | 9.5 | 0.5 |
| 23 | 11 | 11 | 11 | 0 |
| 24 | 10 | 10 | 10 | 0 |
| 25 | 9 | 9 | 8.75 | 0.25 |
| 26 | 9.5 | 9 | 9.25 | -0.25 |
| 27 | 9 | 9 | 8 | 1 |
| 28 | 9 | 9 | 9.25 | -0.25 |
| 29 | 10 | 10 | 10 | 0 |
| 30 | 9.5 | 9 | 8.25 | 0.75 |
| 31 | 11 | 11 | 10.5 | 0.5 |
| 32 | 11.5 | 11 | 8 | 3 |
| 33 | 10 | 10 | 9.75 | 0.25 |
| 34 | 8 | 8 | 8 | 0 |
| 35 | 11 | 11 | 11.25 | -0.25 |
| 36 | 11 | 11 | 11.25 | -0.25 |
| 37 | 10.5 | 10 | 10.5 | -0.5 |
| 38 | 10 | 10 | 10 | 0 |
| 39 | 10.5 | 11 | 10 | 0.5 |
| 40 | 10.5 | 11 | 10.75 | 0.25 |

**Supplementary Table S6:**
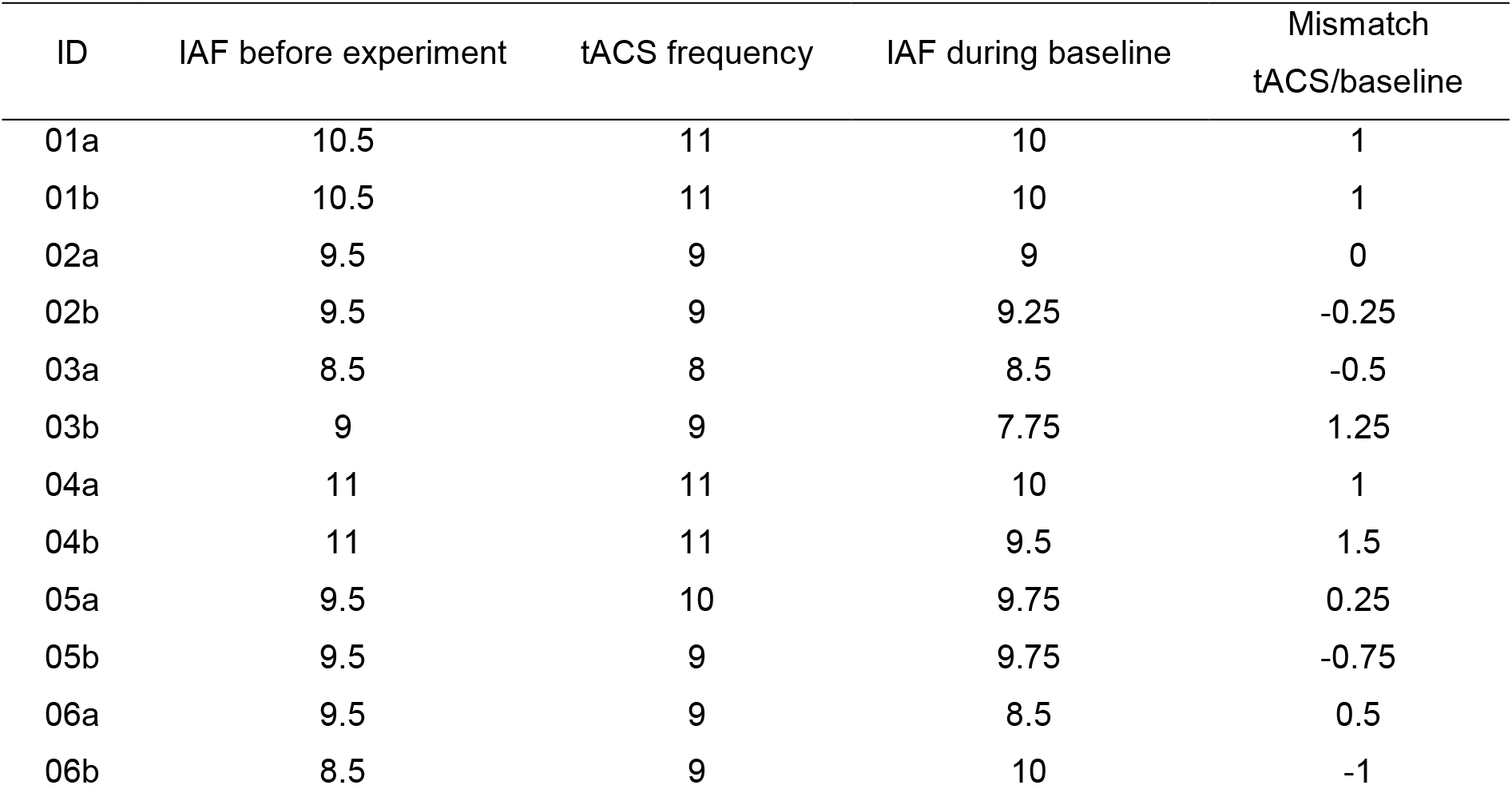
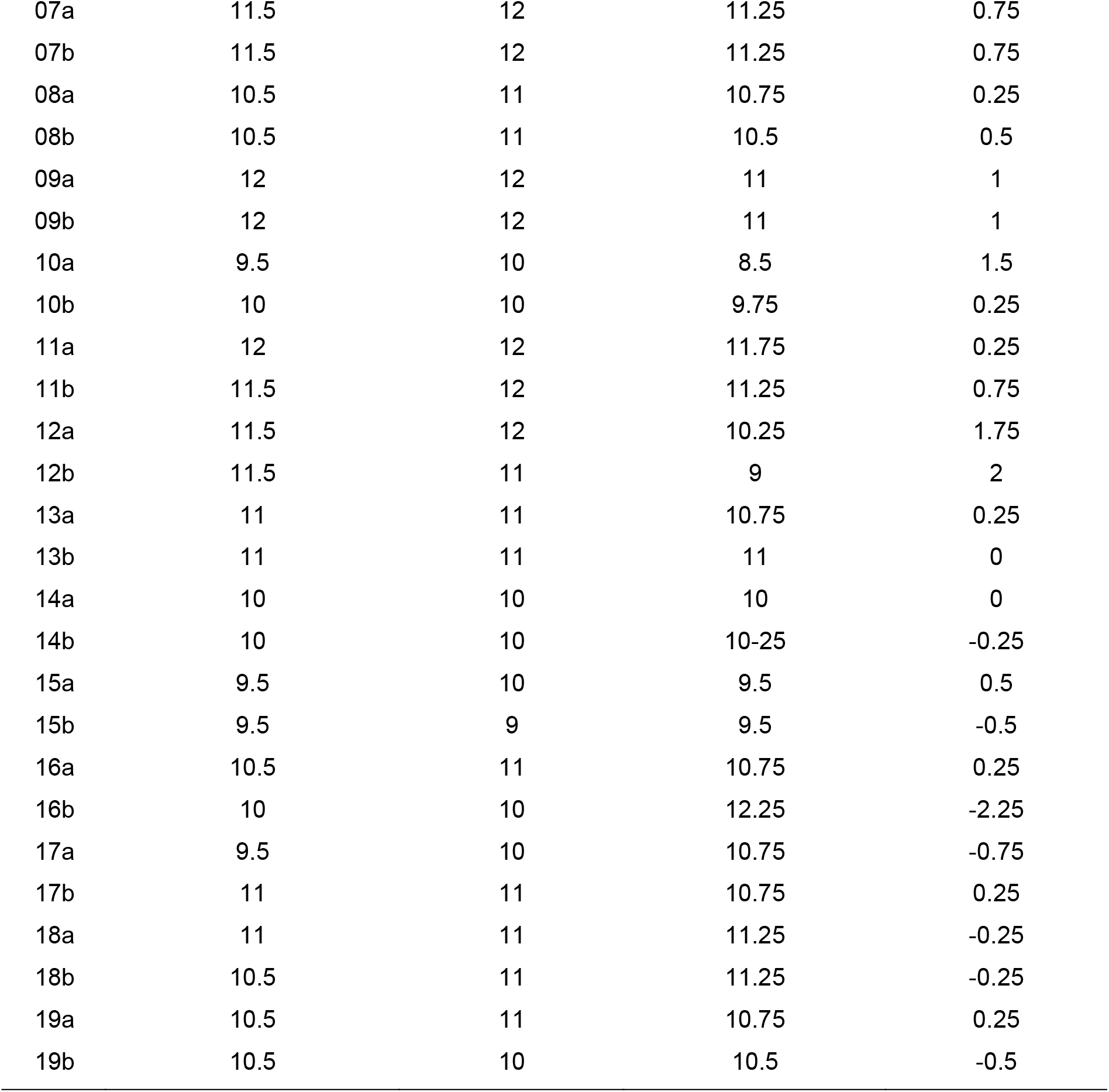
Overview of individual α-frequency measured before the experiment, stimulation frequency during the experiment, IAF measured during the baseline block and the mismatch between the tACS frequency and the IAF during the baseline block of the second experiment.

| ID | IAF before experiment | tACS frequency | IAF during baseline | Mismatch<br>tACS/baseline |
| --- | --- | --- | --- | --- |
| 01a | 10.5 | 11 | 10 | 1 |
| 01b | 10.5 | 11 | 10 | 1 |
| 02a | 9.5 | 9 | 9 | 0 |
| 02b | 9.5 | 9 | 9.25 | -0.25 |
| 03a | 8.5 | 8 | 8.5 | -0.5 |
| 03b | 9 | 9 | 7.75 | 1.25 |
| 04a | 11 | 11 | 10 | 1 |
| 04b | 11 | 11 | 9.5 | 1.5 |
| 05a | 9.5 | 10 | 9.75 | 0.25 |
| 05b | 9.5 | 9 | 9.75 | -0.75 |
| 06a | 9.5 | 9 | 8.5 | 0.5 |
| 06b | 8.5 | 9 | 10 | -1 |
| 07a | 11.5 | 12 | 11.25 | 0.75 |
| 07b | 11.5 | 12 | 11.25 | 0.75 |
| 08a | 10.5 | 11 | 10.75 | 0.25 |
| 08b | 10.5 | 11 | 10.5 | 0.5 |
| 09a | 12 | 12 | 11 | 1 |
| 09b | 12 | 12 | 11 | 1 |
| 10a | 9.5 | 10 | 8.5 | 1.5 |
| 10b | 10 | 10 | 9.75 | 0.25 |
| 11a | 12 | 12 | 11.75 | 0.25 |
| 11b | 11.5 | 12 | 11.25 | 0.75 |
| 12a | 11.5 | 12 | 10.25 | 1.75 |
| 12b | 11.5 | 11 | 9 | 2 |
| 13a | 11 | 11 | 10.75 | 0.25 |
| 13b | 11 | 11 | 11 | 0 |
| 14a | 10 | 10 | 10 | 0 |
| 14b | 10 | 10 | 10-25 | -0.25 |
| 15a | 9.5 | 10 | 9.5 | 0.5 |
| 15b | 9.5 | 9 | 9.5 | -0.5 |
| 16a | 10.5 | 11 | 10.75 | 0.25 |
| 16b | 10 | 10 | 12.25 | -2.25 |
| 17a | 9.5 | 10 | 10.75 | -0.75 |
| 17b | 11 | 11 | 10.75 | 0.25 |
| 18a | 11 | 11 | 11.25 | -0.25 |
| 18b | 10.5 | 11 | 11.25 | -0.25 |
| 19a | 10.5 | 11 | 10.75 | 0.25 |
| 19b | 10.5 | 10 | 10.5 | -0.5 |

## References

1. Filmer, H. L., Dux, P. E. & Mattingley, J. B. Applications of transcranial direct current stimulation for understanding brain function. Trends Neurosci. 37, 742–753 (2014).

2. Herrmann, C. S., Strüber, D., Helfrich, R. F. & Engel, A. K. EEG oscillations: From correlation to causality. Int. J. Psychophysiol. 103, 12–21 (2016).

3. Antal, A. et al. Low intensity transcranial electric stimulation: Safety, ethical, legal regulatory and application guidelines. Clin. Neurophysiol. (2017). doi:10.1016/j.clinph.2017.06.001

4. Bikson, M. et al. Rigor and reproducibility in research with transcranial electrical stimulation: An NIMH-sponsored workshop. Brain Stimul. 11, 465–480 (2018).

5. Kekic, M., Boysen, E., Campbell, I. C. & Schmidt, U. A systematic review of the clinical efficacy of transcranial direct current stimulation (tDCS) in psychiatric disorders. J. Psychiatr. Res. 74, 70–86 (2016).

6. Ahn, S. et al. Targeting reduced neural oscillations in patients with schizophrenia by transcranial alternating current stimulation. Neuroimage 186, 126–136 (2019).

7. Mellin, J. M. et al. Randomized trial of transcranial alternating current stimulation for treatment of auditory hallucinations in schizophrenia. Eur. Psychiatry 51, 25–33 (2018).

8. Bikson, M. et al. Effects of uniform extracellular DC electric fields on excitability in rat hippocampal slices in vitro. J. Physiol. 557, 175–190 (2004).

9. Creutzfeldt, O. D., Fromm, G. H. & Kapp, H. Influence of transcortical d-c currents on cortical neuronal activity. Exp. Neurol. 5, 436–452 (1962).

10. Bindman, L. J., Lippold, O. C. J. & Redfearn, J. W. T. The action of brief polarizing currents on the cerebral cortex of the rat (1) during current flow and (2) in the production of long-lasting after-effects. J. Physiol. 172, 369–382 (1964).

11. Fröhlich, F. & McCormick, D. A. Endogenous Electric Fields May Guide Neocortical Network Activity. Neuron 67, 129–143 (2010).

12. Nitsche, M. a & Paulus, W. Sustained excitability elevations induced by transcranial DC motor cortex stimulation in humans. Neurology 57, 1899–1901 (2001).

13. Kasten, F. H., Dowsett, J. & Herrmann, C. S. Sustained Aftereffect of α-tACS Lasts Up to 70 min after Stimulation. Front. Hum. Neurosci. 10, 245 (2016).

14. Wischnewski, M. et al. NMDA Receptor-Mediated Motor Cortex Plasticity After 20 Hz Transcranial Alternating Current Stimulation. Cereb. Cortex 1–8 (2018). doi:10.1093/cercor/bhy160

15. Nitsche, M. a et al. Pharmacological modulation of cortical excitability shifts induced by transcranial direct current stimulation in humans. J. Physiol. 553, 293–301 (2003).

16. Zaehle, T., Rach, S. & Herrmann, C. S. Transcranial Alternating Current Stimulation Enhances Individual Alpha Activity in Human EEG. PLoS One 5, 13766 (2010).

17. Vossen, A., Gross, J. & Thut, G. Alpha Power Increase After Transcranial Alternating Current Stimulation at Alpha Frequency (α-tACS) Reflects Plastic Changes Rather Than Entrainment. Brain Stimul. 8, 499–508 (2015).

18. Horvath, J. C., Forte, J. D. & Carter, O. Evidence that transcranial direct current stimulation (tDCS) generates little-to-no reliable neurophysiologic effect beyond MEP amplitude modulation in healthy human subjects: A systematic review. Neuropsychologia 66, 213–236 (2015).

19. Horvath, J. C., Forte, J. D. & Carter, O. Quantitative review finds no evidence of cognitive effects in healthy populations from single-session transcranial direct current stimulation (tDCS). Brain Stimul. 8, 535–550 (2015).

20. Veniero, D., Benwell, C. S. Y., Ahrens, M. M. & Thut, G. Inconsistent effects of parietal α-tACS on Pseudoneglect across two experiments: A failed internal replication. Front. Psychol. 8, 1–14 (2017).

21. Fekete, T., Nikolaev, A. R., Knijf, F. De, Zharikova, A. & van Leeuwen, C. Multi-electrode alpha tACS during varying background tasks fails to modulate subsequent alpha power. Front. Neurosci. 12, (2018).

22. Vöröslakos, M. et al. Direct effects of transcranial electric stimulation on brain circuits in rats and humans. Nat. Commun. 9, 483 (2018).

23. Lafon, B. et al. Low frequency transcranial electrical stimulation does not entrain sleep rhythms measured by human intracranial recordings. Nat. Commun. 8, 1–14 (2017).

24. Ridding, M. C. & Ziemann, U. Determinants of the induction of cortical plasticity by non-invasive brain stimulation in healthy subjects. J. Physiol. 588, 2291–304 (2010).

25. Thirugnanasambandam, N. et al. Nicotinergic impact on focal and non-focal neuroplasticity induced by non-invasive brain stimulation in non-smoking humans. Neuropsychopharmacology 36, 879–86 (2011).

26. Grundey, J. et al. Rapid effect of nicotine intake on neuroplasticity in non-smoking humans. Front. Pharmacol. 3 OCT, 1–9 (2012).

27. Krause, B. & Cohen Kadosh, R. Not all brains are created equal: the relevance of individual differences in responsiveness to transcranial electrical stimulation. Front. Syst. Neurosci. 8, 25 (2014).

28. Feurra, M. et al. State-dependent effects of transcranial oscillatory currents on the motor system: what you think matters. J. Neurosci. 33, 17483–9 (2013).

29. Silvanto, J., Muggleton, N. & Walsh, V. State-dependency in brain stimulation studies of perception and cognition. Trends Cogn. Sci. 12, 447–454 (2008).

30. Laakso, I., Tanaka, S., Koyama, S., De Santis, V. & Hirata, A. Inter-subject variability in electric fields of motor cortical tDCS. Brain Stimul. 8, 906–913 (2015).

31. Opitz, A. et al. On the importance of precise electrode placement for targeted transcranial electric stimulation. Neuroimage 181, 560–567 (2018).

32. Neuling, T., Wagner, S., Wolters, C. H., Zaehle, T. & Herrmann, C. S. Finite-element model predicts current density distribution for clinical applications of tDCS and tACS. Front. Psychiatry 3, 1–10 (2012).

33. Thielscher, A., Antunes, A. & Saturnino, G. B. Field modeling for transcranial magnetic stimulation: A useful tool to understand the physiological effects of TMS? Conf. Proc. … Annu. Int. Conf. IEEE Eng. Med. Biol. Soc. IEEE Eng. Med. Biol. Soc. Annu. Conf. 2015, 222–5 (2015).

34. Huang, Y., Datta, A., Bikson, M. & Parra, L. C. Realistic volumetric-approach to simulate transcranial electric stimulation—ROAST—a fully automated open-source pipeline. J.Neural Eng. 16, (2019).

35. Huang, Y. et al. Measurements and models of electric fields in the in vivo human brain during transcranial electric stimulation. Elife 6, 1–27 (2017).

36. Opitz, A. et al. Spatiotemporal structure of intracranial electric fields induced by transcranial electric stimulation in humans and nonhuman primates. Sci. Rep. 6, 1–11 (2016).

37. Neuling, T., Rach, S. & Herrmann, C. S. Orchestrating neuronal networks: sustained after-effects of transcranial alternating current stimulation depend upon brain states. Front. Hum. Neurosci. 7, 161 (2013).

38. Stecher, H. I. & Herrmann, C. S. Absence of Alpha-tACS Aftereffects in Darkness Reveals Importance of Taking Derivations of Stimulation Frequency and Individual Alpha Variability Into Account. Front. Psychol. 9, 1–9 (2018).

39. Benwell, C. S. Y. et al. Frequency and power of human alpha oscillations drift systematically with time-on-task. Neuroimage 192, 101–114 (2019).

40. Haegens, S., Cousijn, H., Wallis, G., Harrison, P. J. & Nobre, A. C. Inter- and intra-individual variability in alpha peak frequency. Neuroimage 92, 46–55 (2014).

41. Gross, J. et al. Dynamic imaging of coherent sources: Studying neural interactions in the human brain. Proc. Natl. Acad. Sci. 98, 694–699 (2001).

42. Schutter, D. J. L. G. Cutaneous retinal activation and neural entrainment in transcranial alternating current stimulation: A systematic review. Neuroimage 140, 83–88 (2016).

43. Asamoah, B., Khatoun, A. & Mc Laughlin, M. tACS motor system effects can be caused by transcutaneous stimulation of peripheral nerves. Nat. Commun. 10, 266 (2019).

44. Wagner, S., Burger, M. & Wolters, C. H. An Optimization Approach for Well-Targeted Transcranial Direct Current Stimulation. SIAM J. Appl. Math. 76, 2154–2174 (2016).

45. Bergmann, T. O., Karabanov, A., Hartwigsen, G., Thielscher, A. & Siebner, H. R. Combining non-invasive transcranial brain stimulation with neuroimaging and electrophysiology: Current approaches and future perspectives. Neuroimage 140, 4–19 (2016).

46. Kar, K. & Krekelberg, B. Transcranial electrical stimulation over visual cortex evokes phosphenes with a retinal origin. J. Neurophysiol. 108, 2173–2178 (2012).

47. Antal, A. & Herrmann, C. S. Transcranial Alternating Current and Random Noise Stimulation: Possible Mechanisms. Neural Plast. 2016, 1–12 (2016).

48. Reato, D., Rahman, A., Bikson, M. & Parra, L. C. Low-Intensity Electrical Stimulation Affects Network Dynamics by Modulating Population Rate and Spike Timing. J. Neurosci. 30, 15067–15079 (2010).

49. Ozen, S. et al. Transcranial Electric Stimulation Entrains Cortical Neuronal Populations in Rats. J. Neurosci. 30, 11476–11485 (2010).

50. Ali, M. M., Sellers, K. K. & Frohlich, F. Transcranial Alternating Current Stimulation Modulates Large-Scale Cortical Network Activity by Network Resonance. J. Neurosci. 33, 11262–11275 (2013).

51. Siegel, M., Donner, T. H. & Engel, A. K. Spectral fingerprints of large-scale neuronal interactions. Nat. Rev. Neurosci. 13, 121–134 (2012).

52. Daniel, R. S. Alpha and theta EEG in vigilance. Percept. Mot. Skills 25, 697–703 (1967).

53. Oken, B. S., Salinsky, M. C. & Elsas, S. M. Vigilance, alertness, or sustained attention: physiological basis and measurement. Clin. Neurophysiol. 117, 1885–1901 (2006).

54. Boksem, M. A. S., Meijman, T. F. & Lorist, M. M. Effects of mental fatigue on attention: An ERP study. Cogn. Brain Res. 25, 107–116 (2005).

55. Cajochen, C., Brunner, D. P., Krauchi, K., Graw, P. & Wirz-Justice, A. Power Density in Theta/Alpha Frequencies of the Waking EEG Progressively Increases During Sustained Wakefulness. Sleep 18, 890–894 (1995).

56. Rahman, A. et al. Cellular effects of acute direct current stimulation: somatic and synaptic terminal effects. J. Physiol. 591, 2563–2578 (2013).

57. Oldfield, R. C. C. The assessment and analysis of handedness: The Edinburgh inventory. Neuropsychologia 9, 97–113 (1971).

58. Noury, N., Hipp, J. F. & Siegel, M. Physiological processes non-linearly affect electrophysiological recordings during transcranial electric stimulation. Neuroimage 140, 99–109 (2016).

59. Noury, N. & Siegel, M. Phase properties of transcranial electrical stimulation artifacts in electrophysiological recordings. Neuroimage 158, 406–416 (2017).

60. Kasten, F. H., Maess, B. & Herrmann, C. S. Facilitated Event-Related Power Modulations during Transcranial Alternating Current Stimulation (tACS) Revealed by Concurrent tACS-MEG. eneuro 5, ENEURO.0069-18.2018 (2018).

61. Kasten, F. H., Negahbani, E., Fröhlich, F. & Herrmann, C. S. Non-linear transfer characteristics of stimulation and recording hardware account for spurious low-frequency artifacts during amplitude modulated transcranial alternating current stimulation (AM-tACS). Neuroimage 179, 134–143 (2018).

62. Brunoni, A. R. et al. A systematic review on reporting and assessment of adverse effects associated with transcranial direct current stimulation. Int. J. Neuropsychopharmacol. 14, 1133–1145 (2011).

63. Stecher, H. I. et al. Ten Minutes of α-tACS and Ambient Illumination Independently Modulate EEG α-Power. Front. Hum. Neurosci. 11, 1–10 (2017).

64. Kleiner, M., Brainard, D. & Pelli, D. What’s new in Psychtoolbox-3? in Perception 36 ECVP Abstract Supplement (2007). doi:10.1068/v070821

65. Oostenveld, R., Fries, P., Maris, E. & Schoffelen, J. M. FieldTrip: Open source software for advanced analysis of MEG, EEG, and invasive electrophysiological data. Comput. Intell. Neurosci. 2011, 1–9 (2011).

66. Taulu, S. & Simola, J. Spatiotemporal signal space separation method for rejecting nearby interference in MEG measurements. Phys. Med. Biol. 51, 1759–1768 (2006).

67. Taulu, S., Simola, J. & Kajola, M. Applications of the signal space separation method. IEEE Trans. Signal Process. 53, 3359–3372 (2005).

68. Nenonen, J. et al. Validation of head movement correction and spatiotemporal signal space separation in magnetoencephalography. Clin. Neurophysiol. 123, 2180–2191 (2012).

69. Medvedovsky, M., Taulu, S., Bikmullina, R., Ahonen, A. & Paetau, R. Fine tuning the correlation limit of spatio-temporal signal space separation for magnetoencephalography. J. Neurosci. Methods 177, 203–211 (2009).

70. Debener, S., Thorne, J., Schneider, T. R. & Viola, F. C. Using ICA for the Analysis of Multi-Channel EEG Data. in Simultaneous EEG and fMRI : Recording, Analysis, and Application (2010). doi:10.1093/acprof:oso/9780195372731.003.0008

71. Nolte, G. The magnetic lead field theorem in the quasi-static approximation and its use for magnetoencephalography forward calculation in realistic volume conductors. Phys. Med. Biol. 48, 3637–3652 (2003).

